# Prolonged Cell Cycle Arrest in Response to DNA damage in Yeast Requires the Maintenance of DNA Damage Signaling and the Spindle Assembly Checkpoint

**DOI:** 10.1101/2023.05.15.540538

**Authors:** Felix Y. Zhou, David P. Waterman, Marissa Ashton, Suhaily Caban-Penix, Gonen Memisoglu, Vinay V. Eapen, James E. Haber

## Abstract

Cells evoke the DNA damage checkpoint (DDC) to inhibit mitosis in the presence of DNA double-strand breaks (DSBs) to allow more time for DNA repair. In budding yeast, a single irreparable DSB is sufficient to activate the DDC and induce cell cycle arrest prior to anaphase for about 12 to 15 hours, after which cells “adapt” to the damage by extinguishing the DDC and resuming the cell cycle. While activation of the DNA damage-dependent cell cycle arrest is well-understood, how it is maintained remains unclear. To address this, we conditionally depleted key DDC proteins after the DDC was fully activated and monitored changes in the maintenance of cell cycle arrest. Degradation of Ddc2^ATRIP^, Rad9, Rad24, or Rad53^CHK2^ results in premature resumption of the cell cycle, indicating that these DDC factors are required both to establish and to maintain the arrest. Dun1 is required for establishment, but not maintenance of arrest, whereas Chk1 is required for prolonged maintenance but not for initial establishment of the mitotic arrest. When the cells are challenged with 2 persistent DSBs, they remain permanently arrested. This permanent arrest is initially dependent on the continuous presence of Ddc2, Rad9, and Rad53; however, after 15 hours these proteins become dispensable. Instead, the continued mitotic arrest is sustained by spindle-assembly checkpoint (SAC) proteins Mad1, Mad2, and Bub2 but not by Bub2’s binding partner Bfa1. These data suggest that prolonged cell cycle arrest in response to 2 DSBs is achieved by a handoff from the DDC to specific components of the SAC. Furthermore, the establishment and maintenance of DNA damage-induced cell cycle arrest requires overlapping but different sets of factors.

## Introduction

DNA double-strand breaks (DSBs) are one of the most deleterious forms of DNA damage (Mehta and Haber 2014). In response to DSBs, cells evoke the DNA damage checkpoint (DDC) to halt the metaphase to anaphase transition (known as the G_2_/M checkpoint). Activation of DDC gives cells an extended opportunity to repair DSBs and therefore, prevents the inheritance of broken chromosomes, which can lead to aneuploidy, chromosome aberrations and genome instability (Waterman, Haber, and Smolka 2020).

In budding yeast, a single irreparable DSB is sufficient to trigger the DDC through the activation of Mec1, a PI3K-like kinase and homolog of the mammalian ATR (Mantiero et al. 2007; Pellicioli et al. 2001). Mec1 activation depends on 5’ to 3’ resection of the DSB ends that exposes single-stranded DNA (ssDNA) which is rapidly coated by the ssDNA binding protein, RPA (reviewed by (Marechal and Zou 2015)). As resection proceeds, the PCNA-related 9-1-1 clamp, made up of Ddc1, Rad17, and Mec3, is loaded at the resected ss/dsDNA junction by the Rad24-Rfc2-5 clamp loader (Ellison and Stillman 2003; Majka et al. 2006). Mec1 is recruited to DSB sites via its obligate binding partner, Ddc2^ATRIP^ interacting with RPA-bound ssDNA (Dubrana et al. 2007; Zou and Elledge 2003). Following its localization to DSBs, Mec1’s kinase activity is stimulated by Dbp11, Dna2 and the Ddc1 subunit of the 9-1-1 clamp (Navadgi-Patil and Burgers 2009; Melo, Cohen, and Toczyski 2001). Impairing Mec1’s kinase activity by the PI3K-like kinase inhibitor caffeine, by using temperature-sensitive Mec1 mutants or by degradation of Mec1’s binding partner Ddc2, rapidly extinguishes checkpoint signaling (Pellicioli et al. 2001; Tsabar et al. 2015; Vaze et al. 2002), illustrating that continual Mec1 activity is needed to activate and sustain DDC. In contrast to Mec1, yeast’s other PI3K-like kinase, Tel1, the homolog of mammalian ATM, is dispensable for DDC activation and maintenance, as *TEL1* deletion only shortens damage-induced cell cycle arrest by a few hours (Dubrana et al. 2007).

Following the induction of a DSB, numerous proteins are phosphorylated either directly by Mec1 or by the downstream effector kinases Rad53 and Chk1 (human CHK2 and CHK1), which are themselves Mec1 substrates (Lanz, Dibitetto, and Smolka 2019; Smolka et al. 2007). Mec1 and Tel1 substrates also include histone H2A-S129, called γ-H2AX, which spreads on both sides of the DSB via two distinct mechanisms (Rogakou et al. 1998; Shroff et al. 2004). γ-H2AX then recruits the scaffold protein Rad9, which brings the effector kinase Rad53 in close proximity to Mec1 for activation (Durocher et al. 1999; Emili 1998; Schwartz et al. 2002). Activated Rad53 then amplifies the DDC signal through autophosphorylation in *trans*, also stimulating the transcription regulator Dun1 kinase, while restraining the degradation of Pds1 (securin) to inhibit mitosis (Chen, Smolka, and Zhou 2007; Fiorani et al. 2008; Pellicioli et al. 1999; Usui and Petrini 2007; Yam et al. 2020).

In addition to the DDC, unattached kinetochores can induce cell cycle arrest through the activation of spindle assembly checkpoint (SAC) (reviewed by (Musacchio 2015)). Several studies have suggested a crosstalk between the SAC and the DDC. For example, deletion of SAC components *MAD1* or *MAD2* shorten the cell cycle arrest in response to DNA damaging agents in the absence of DDC genes *RAD9* and *RAD24* (Garber and Rine 2002; Kim and Burke 2008). Furthermore, *MAD1, MAD2* or *BUB1* mutants arrest for less time than wild-type cells following the induction of a single persistent DSB (Dotiwala et al. 2010). In mouse oocytes, inhibition of the SAC overrides the activation of DDC-mediated metaphase arrest during the first meiotic division (Marangos et al. 2015). The mitotic exit network (MEN) is another signaling cascade activated during anaphase to promote cell cycle re-entry (Matellan and Monje-Casas 2020); therefore, defects in MEN lead to mitotic arrest in late anaphase (Geymonat et al. 2002; Bardin, Visintin, and Amon 2000; Shirayama, Matsui, and Toh 1994). In addition to SAC, MEN also communicates with the DDC in response to DNA damage. For instance, a key regulator of MEN, the heterodimer Bub2/Bfa1 complex, is modified in a Rad53 and Dun1-dependent manner following damage (Hu et al. 2001). Supporting the idea of crosstalk between MEN and DDC, our lab had shown that deletion of *BUB2* shortened the duration of the arrest in response to a single unrepaired DSB (Dotiwala et al. 2010).

Here, we present new mechanistic insights into the *maintenance* of the cell cycle arrest following DNA damage by employing the auxin-inducible degron (AID) strategy to conditionally deplete DDC and SAC proteins. An advantage of the AID system, compared to null or temperature-sensitive mutants, is that AID-tagged proteins retain wild-type function until the addition of the plant hormone auxin (indole-3-acetic acid, IAA), which triggers rapid degradation of AID-tagged proteins in the presence of the TIR1 E3 ubiquitin ligase (Morawska and Ulrich 2013; Nishimura et al. 2009). To investigate how conditional depletion of DDC or SAC proteins alter the maintenance of cell cycle arrest, we engineered a yeast strain that permanently arrests due to the presence of 2 persistent DNA breaks. We find that the maintenance of the cell cycle arrest requires constant presence of some, but not all, checkpoint activation proteins. Surprisingly, we find that the DDC proteins which are essential to induce the cell cycle arrest and sustain it at the early stages of the arrest become dispensable nearly 15 hours after DNA damage induction. Conversely, SAC proteins are dispensable for the establishment and the initial steps of the cell cycle arrest but become essential at later stages of the DNA damage dependent cell cycle arrest. Based on these findings, we posit that prolonged cell cycle arrest in response to DNA damage is sustained by both SAC and DDC; however, each checkpoint sustains the arrest at different stages.

## Results

### Measuring DNA Damage Checkpoint Arrest and Maintenance

To study the role of DDC initiation proteins in the *maintenance* of the cell cycle arrest, we utilized the well-characterized strain JKM179 (Lee et al. 1998), in which the site-specific HO endonuclease is expressed from a *GAL1-10* promoter (*GAL-HO*) to induce a single DSB within the *MAT* locus on chromosome III (referred to as the 1-DSB strain). In this 1-DSB strain, we inserted an additional HO cleavage site 52 kb from the centromere on chromosome IV to induce another DSB (referred to as the 2-DSB strain) (Kim et al. 2007; Lee et al. 1998). At both loci, HO-mediated cleavage after galactose induction is nearly complete within 30-45 minutes (Lee et al. 2014). In both strains, we also deleted the *HML* and *HMR* donors to prevent repair by homologous recombination. With continuous *HO* expression, nonhomologous end-joining occurs in only 0.2% of these cells (Moore and Haber 1996) thus both DSBs are essentially irreparable.

Following the induction of two DSBs, we monitored cell cycle arrest in 4 ways: (1) with an adaptation time-course assay where we micromanipulated individual G_1_ cells on agar plates and scored the percentage of cells that are able to re-enter mitosis (Lee et al. 1998), (2) by monitoring the percentage of G_2_/M-arrested cells in liquid culture based on cell morphology (**Figure 1A**), (3) by monitoring nuclear division by DAPI staining of nuclei, and (4) by assaying Rad53 phosphorylation by western blot (Pellicioli et al. 2001).

**Figure 1.**
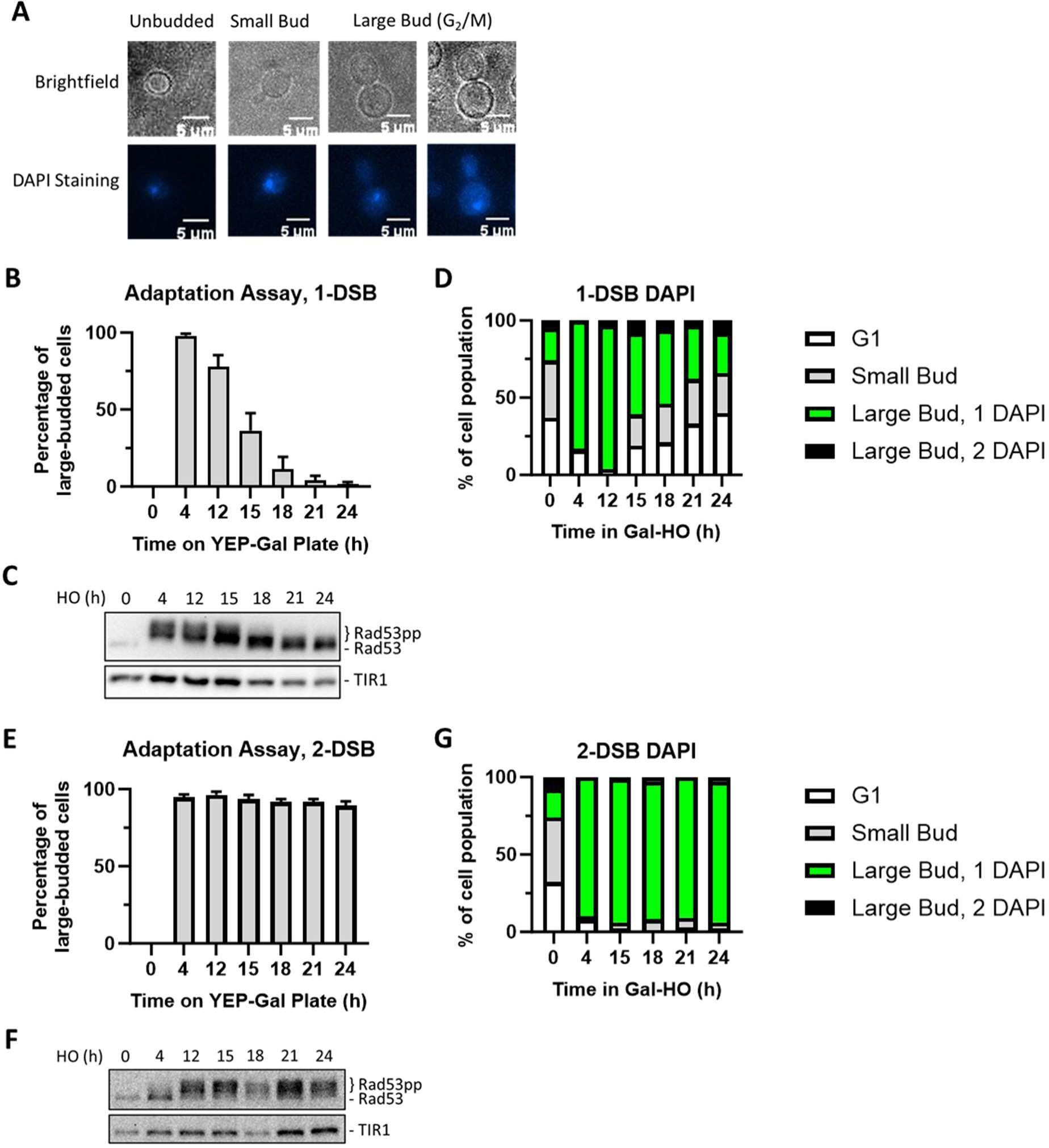
Measuring checkpoint arrest in a 1-DSB and 2-DSB strains. (A) Morphological categories of budding yeast cells using brightfield microscopy and DAPI staining were used to determine G_2_/M arrest. Cells that arrest at G2/M shift towards a large bud state. G_2_/M arrested cells that progress into anaphase. (B) Adaptation assay with 1 DSB strain on a YEP-Gal plate. G_2_/M arrest was determined based on cell morphology as shown in Fig. 1A. Data are shown from 3 independent experiments, error bars represent standard error of the mean (SEM). (C) Profile of DAPI stained cells in a 1-DSB strain after DNA damage induction in liquid culture. Cells were grouped based on cell morphology and DAPI staining profiles, as explained below the graphs. (D) Rad53 phosphorylation kinetics in 1 DSB strain by western blotting. Samples collected after the induction of DNA damage during the time course experiment and blotted with α-Rad53 to monitor DDC signaling. α-Rad53 can both detect unphosphorylated and hyperphosphorylated Rad53 species. TIR1-Myc was detected with α-Myc and serves as a loading control. (E) Same as (B) for a 2-DSB strain. (F) Same as (C) with a 2-DSB strain. (G) Same as (D) with a 2-DSB strain.

In both 1-DSB and 2-DSB strains, 4 h after induction of DNA damage, >90% of cells arrested at G_2_/M as determined by an adaptation assay (**Figure 1B, 1E)** and DAPI staining **(Figure 1D and 1G).** In agreement, western blot analysis showed that Rad53 was hyperphosphorylated (**Figure 1C and 1F**), demonstrating that DDC was fully activated in both these strains after DNA damage. By 12 to 15 h after the induction of a single persistent DSB, most 1-DSB cells adapted; that is, they escaped the G_2_/M arrest and re-entered mitosis (**Figure 1B**). The timing of Rad53 dephosphorylation following the induction of a single irreparable DSB correlated with the timing of adaptation and escape from the G_2_/M arrest (**Figure 1C**), as previously shown (Pellicioli et al. 2001) In contrast, in the 2-DSB strain, over 90% of cells remained permanently arrested in G_2_/M with persistently hyper-phosphorylated Rad53 throughout the 24 h time-course (**Figure 1E-G**). We leveraged this permanent cell cycle arrest observed in the 2-DSB strain to study how the DNA damage-induced cell cycle arrest is maintained once it had been established.

### Analysis of Checkpoint Factors Required for Checkpoint Maintenance

We used the auxin-inducible degron (AID) system (Nishimura et al. 2009) to conditionally deplete DDC proteins after G_2_/M arrest had been established to study the maintenance of cell cycle arrest. To this end, we appended an AID tag with 9 copies of the c-MYC epitope tag to the C-terminus of Ddc2, Rad9, Rad24, and Rad53, which are all components of the Mec1 signaling cascade (Memisoglu et al. 2019; Sweeney et al. 2005; de la Torre-Ruiz, Green, and Lowndes 1998). Hereafter, all AID-tagged proteins will be designated simply as -AID, e.g., Ddc2-9xMyc-AID as Ddc2-AID.

AID-tagging of DDC proteins did not alter the establishment of G_2_/M arrest; however, *RAD9-AID*, *RAD24-AID*, and *RAD53-AID* strains are hypomorphic and adapted 24 h after DNA damage in the absence of IAA, while a 2-DSB wild-type counterpart cells remained fully arrested (**Figure S1A**). This premature escape from the cell cycle arrest was dependent on the presence of TIR1 (**Figure S1B**) and is likely due to low levels of IAA as a natural intermediate in amino acid biosynthesis (Rao et al. 2010). IAA treatment prior to the induction of 2 DSBs in a control strain, which does not contain any AID-tagged proteins, did not alter the prolonged cell cycle arrest following DNA damage (**Figure S2A**), illustrating that IAA treatment by itself does not alter response to DNA damage. However, rapid degradation of Ddc2-AID, Rad9-AID, Rad24-AID and Rad53-AID by IAA treatment 2 h prior to the induction of DSBs largely prevented cell cycle arrest as well as DDC signaling, evident from the absence of detectible Rad53 hyperphosphorylation (**Figure S2B-E**). These results underlie the importance of Ddc2, Rad9, Rad24 and Rad53 in initiating DDC signaling and cell cycle arrest in response to DNA damage, agreeing with previous reports (Pellicioli et al. 2001; Paciotti et al. 2000; Emili 1998; de la Torre-Ruiz, Green, and Lowndes 1998).

To test whether the DDC proteins are required for *maintenance* of the cell cycle arrest following DNA damage, we employed the AID tagged strains with 2 DSBs and depleted the AID proteins 4 h *after* inducing DSBs. In the absence of IAA, *DDC2-AID*, *RAD9-AID*, or *RAD24-AID* strains all activated the DDC signaling 4 h after DSB induction, with 89% - 99% of cells arrested in G_2_/M (**Figure 2A-C**), demonstrating that AID tagging of these proteins did not impair their function. Within 1 h after IAA treatment, Ddc2-AID, Rad9-AID, or Rad24-AID were all rapidly depleted, which caused a gradual Rad53 dephosphorylation, as detected by western blotting (**Figure 2A-C**). Moreover, agreeing with the loss of Rad53 phosphorylation, IAA treatment of DDC-AID strains triggered release from G_2_/M arrest, while the untreated control cells remained fully arrested. DAPI staining of *DDC2-AID* cells after IAA treatment revealed the accumulation of large-budded cells with 2 distinct DAPI signals, indicating that cells started to progress into anaphase following Ddc2 depletion (**Figure S3A**). These findings illustrate that the upstream DDC factors Ddc2, Rad9 and Rad24 are essential for initiating and sustaining the cell cycle arrest in response to DNA damage.

**Figure 2.**
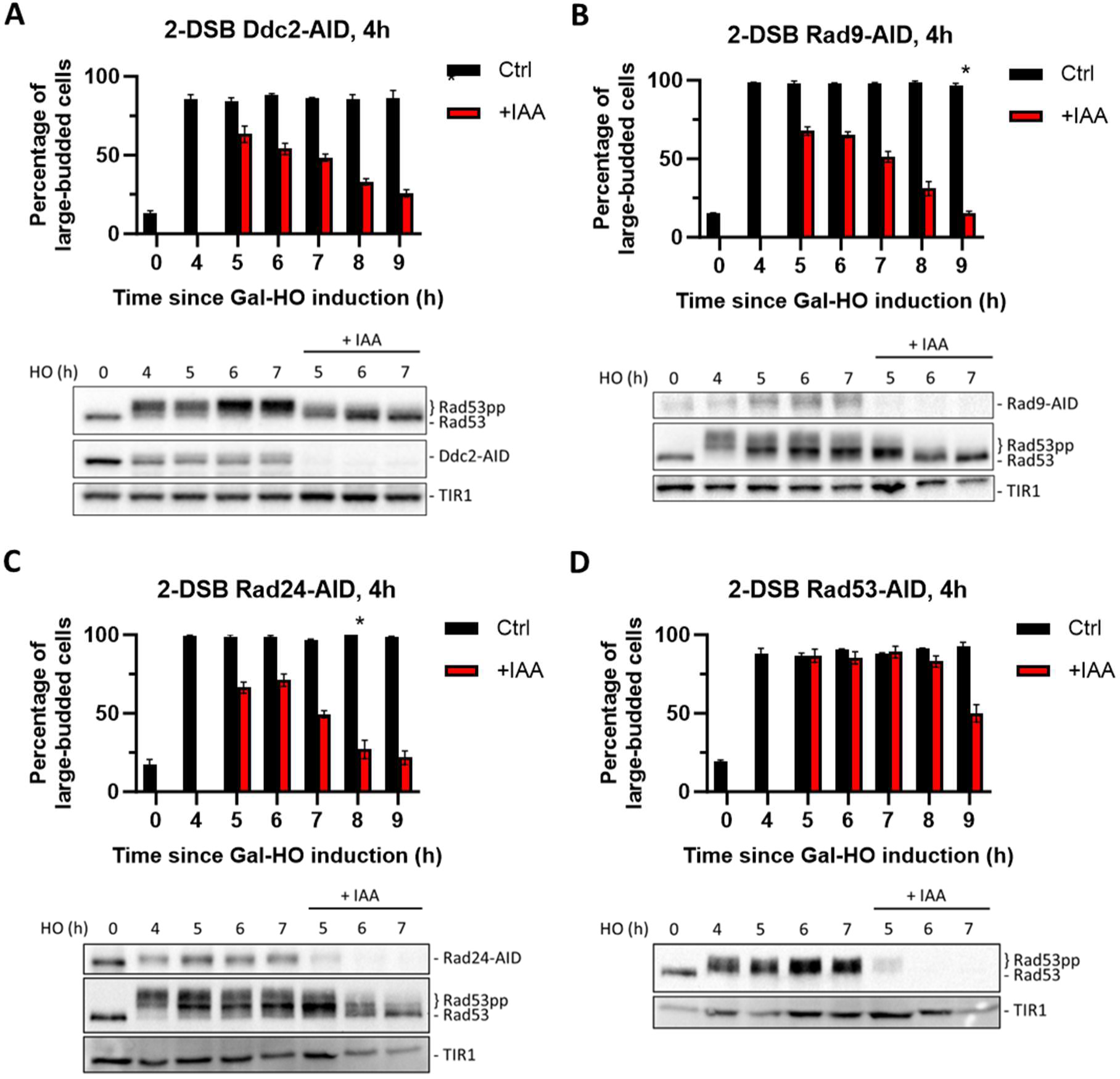
Checkpoint maintenance requires Ddc2, Rad9, Rad24, and Rad53 activity. (A) Above, percentage of G_2_/M arrested cells in a 2-DSB *DDC2-AID* strain after DNA damage induction in a liquid culture. Cultures were split 4 h after galactose treatment to induce DNA damage by GAL::HO and treated either with auxin (+IAA) (1mM) or with ethanol (Ctrl). Data are shown from 3 independent experiments with error bars representing standard error of the mean (SEM). The asterisk marks the time point when the percentage of large-budded G_2_/M cells returned to pre-damage levels. Below, western blots ran with samples collected at various time points during the same time course, probed with α-Rad53, to determine DDC status, and α-Myc, to determine Ddc2-AID-Myc protein abundance and TIR1-Myc as a loading control. (B) Same as (A) for 2-DSB *RAD9-AID*. (C) Same as (A) for 2-DSB *RAD24-AID*. (D) Same as (A) for 2-DSB *RAD53-AID*.

Compared to *DDC2-AID*, *RAD9-AID* or *RAD24-AID*, we found that *RAD53-AID* cells maintained G_2_/M arrest for an additional 4 h after complete depletion of Rad53 (**Figure 2D**). In contrast to Ddc2-AID depletion, cell cycle analysis by DAPI staining showed that Rad53 depletion led to a more gradual transition into late anaphase (**Figure S3B**). This delay after the conditional depletion of Rad53 could be due to continued signaling from downstream targets activated by Rad53 kinase, such as Dun1, or from other targets of the Mec1 kinase, downstream of Ddc2, Rad9 and Rad24.

### Chk1 Sustains Checkpoint Signaling in the Absence of Ddc2, Rad9, Rad24, or Rad53

Rad53 and Chk1 kinases both contribute to the maintenance of the cell cycle arrest after DNA damage (Dotiwala et al. 2007; Pellicioli et al. 2001). Agreeing with previously published reports (Sanchez et al. 1999), we find that Chk1 is involved in maintaining the permanent arrest following the induction of 2 DSBs. Deletion of *CHK1* did not impair the induction of cell cycle arrest **(Figure 3A)**; however, it inhibited the permanent cell cycle arrest, as >95% of *chk1*Δ cells adapted by 24 h (**Figure 3B**). We then asked whether the delay in cell cycle re-entry observed when Rad53 was degraded was due to Chk1’s independent role in maintaining arrest. To test this, we induced 2 DSBs in *RAD53-AID chk1Δ* cells for 4 h and then added IAA to deplete Rad53. Compared to the depletion of Rad53-AID alone **(Figure 2D)**, the depletion of Rad53-AID in the absence of *CHK1* led to a significant decrease in the number of G_2_/M-arrested cells within 1 h of IAA treatment (**Figure 3C).** These results suggest that Chk1 functions in conjunction with Rad53 to sustain cell cycle arrest in response to DNA damage.

**Figure 3.**
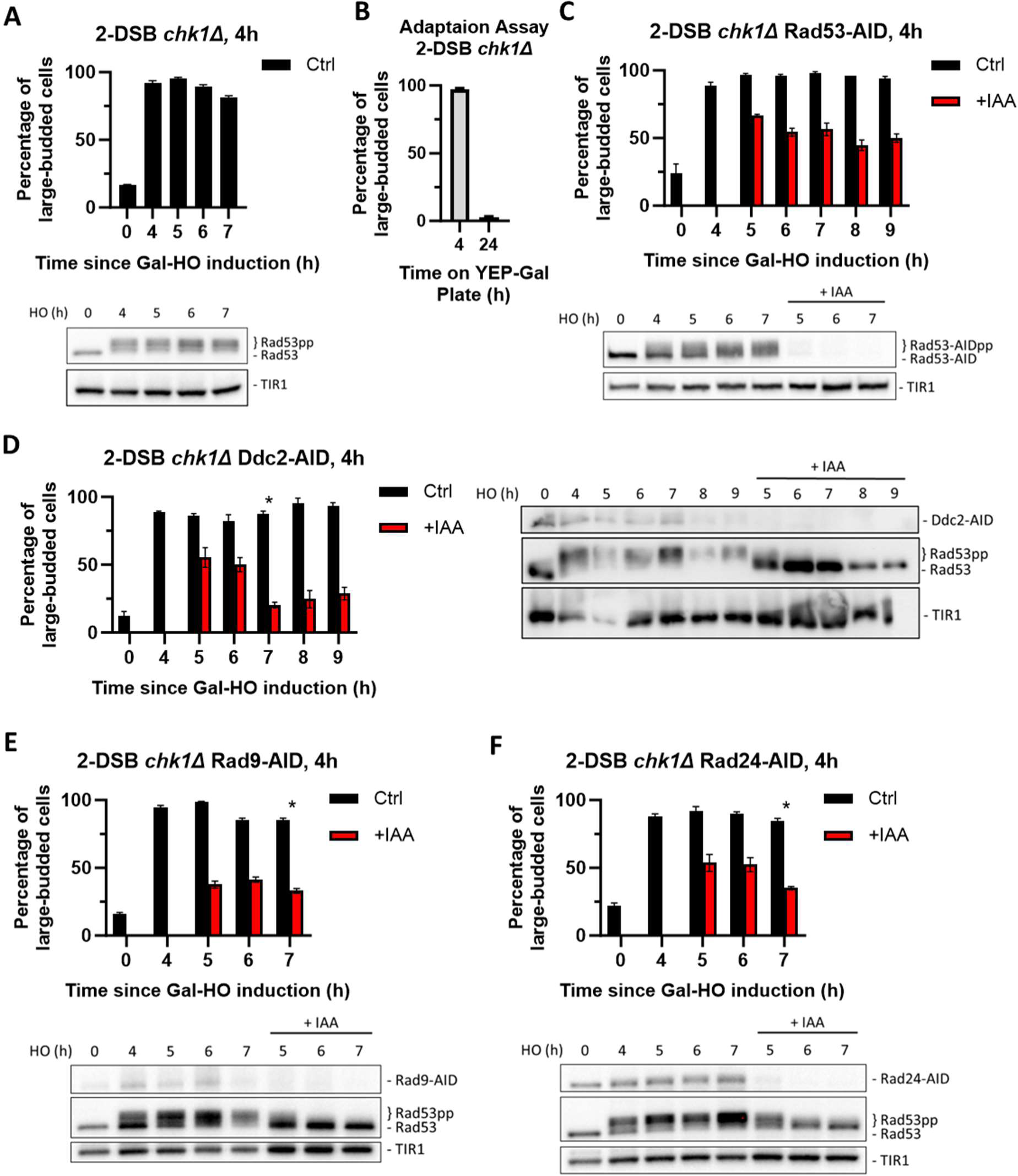
Chk1 is dispensable for activation of the cell cycle arrest, but essential for its maintenance. (A) Percent G_2_/M cells in a 2-DSB *chk1Δ* strain following DNA damage. Data are shown from 3 independent experiments with error bars representing the standard error of the mean (SEM). Western blot probed with α-Rad53 to determine the status of DDC and α-Myc to determine TIR1-Myc protein abundance. (B) Adaptation assay with 2-DSB *chk1*Δ strain. (C) Percentage of G_2_/M arrested cells a 2-DSB *chk1Δ RAD53-AID* strain after DNA damage. Cultures were split 4 h after DSB induction and treated with 1 mM auxin (+IAA) or with ethanol (Ctrl). Data are shown from 3 independent experiments with error bars representing the standard error of the mean (SEM). Western blot probed with α-Myc for Rad53-AID and TIR1-Myc as a loading control. (D) Same as (C) for 2-DSB *chk1Δ DDC2-AID*. Western blot probed with α-Rad53 and α-Myc. α-Rad53 shows both an unphosphorylated protein and multiple phosphorylated species. α-Myc shows Ddc2-AID degradation and TIR1-Myc as a loading control. The asterisk shows when the percentage of large-budded cells returned to pre-damage levels. (E) Same as (D) for 2-DSB *chk1Δ RAD9-AID*. (F) Same as (D) for 2-DSB *chk1Δ RAD24-AID*.

To explore further how Chk1 signaling contributes to the maintenance of DDC-dependent cell cycle arrest, we depleted DDC factors Ddc2-AID, Rad9-AID, or Rad24-AID in *chk1Δ* cells 4 h after the induction of DSBs. Depletion of these upstream factors in the absence of *CHK1* led to a more rapid release from the cell cycle arrest (**Figure 3D-F**) compared to the depletion of DDC factors alone **(Figure 2A-C**). Collectively, these findings demonstrate that Chk1 plays a key role in maintaining cell cycle arrest in response to DNA damage.

Tel1 is thought to play a minor role in response to DSBs, as the establishment of DSB-induced cell cycle arrest normally depends entirely on Mec1 (Dotiwala et al. 2010). However, previous studies have shown that, in addition to Mec1, Tel1 can also target Chk1 for phosphorylation (Limbo et al. 2011; Sanchez et al. 1999). Additionally, when the initial 5’ to 3’ end resection of DSB ends is impaired, Tel1 alone can activate the DDC (Usui and Petrini 2007). To study how Tel1 contributes to the maintenance of the permanent cell cycle arrest, we deleted *TEL1* in 2-DSB strain. Unlike *chk1*Δ with 2 DSBs, a *TEL1* deletion did not affect either the establishment of the DDC nor the maintenance of checkpoint arrest up to 24 h (**Figure S4**), agreeing with previously published results (Dubrana et al. 2007). Taken together, these data illustrate that DNA damage-dependent cell cycle arrest is initiated by Mec1 branch of the DDC via Ddc2, Rad9 and Rad24, and the cell cycle arrest is largely sustained by the downstream kinases Rad53 and Chk1, with minor contributions from other downstream targets of DDC.

### Dun1 is Required for the Initiation but not for the Maintenance of Cell Cycle Arrest

Our findings show that a small number of cells remain arrested in the absence of *CHK1* when Rad53 is depleted. We posited that Rad53 could modulate the expression of other DDC factors which, in turn, sustain the cell cycle arrest in the absence of Rad53 and Chk1. One candidate protein is Dun1, a Rad53-activated protein kinase that regulates transcription in response to DNA damage (Chen, Smolka, and Zhou 2007; Yam et al. 2020; Zhou and Elledge 1993). Deleting *DUN1* significantly impaired checkpoint activation: compared to the wild-type control strain, only 60% of *DUN1* cells arrested in G_2_/M 4 h after the DSB induction, and only 25% remained in G_2_/M arrest at 7 h (**Figure 4A**). Additionally, depletion of Dun1-AID 4 h after damage induction did not cause a significant change in the proportion of G_2_/M arrested in these otherwise wild type cells, nor did it affect Rad53 phosphorylation (**Figure 4B**); however, deleting *CHK1* triggered an exit from checkpoint arrest following the depletion of Dun1-AID 4 h after DSB induction (**Figure 4C**) further demonstrating the role of Chk1 in checkpoint maintenance. These results suggest that Dun1 is required for the initiation of DDC and concomitant cell cycle arrest but is dispensable for maintenance.

**Figure 4.**
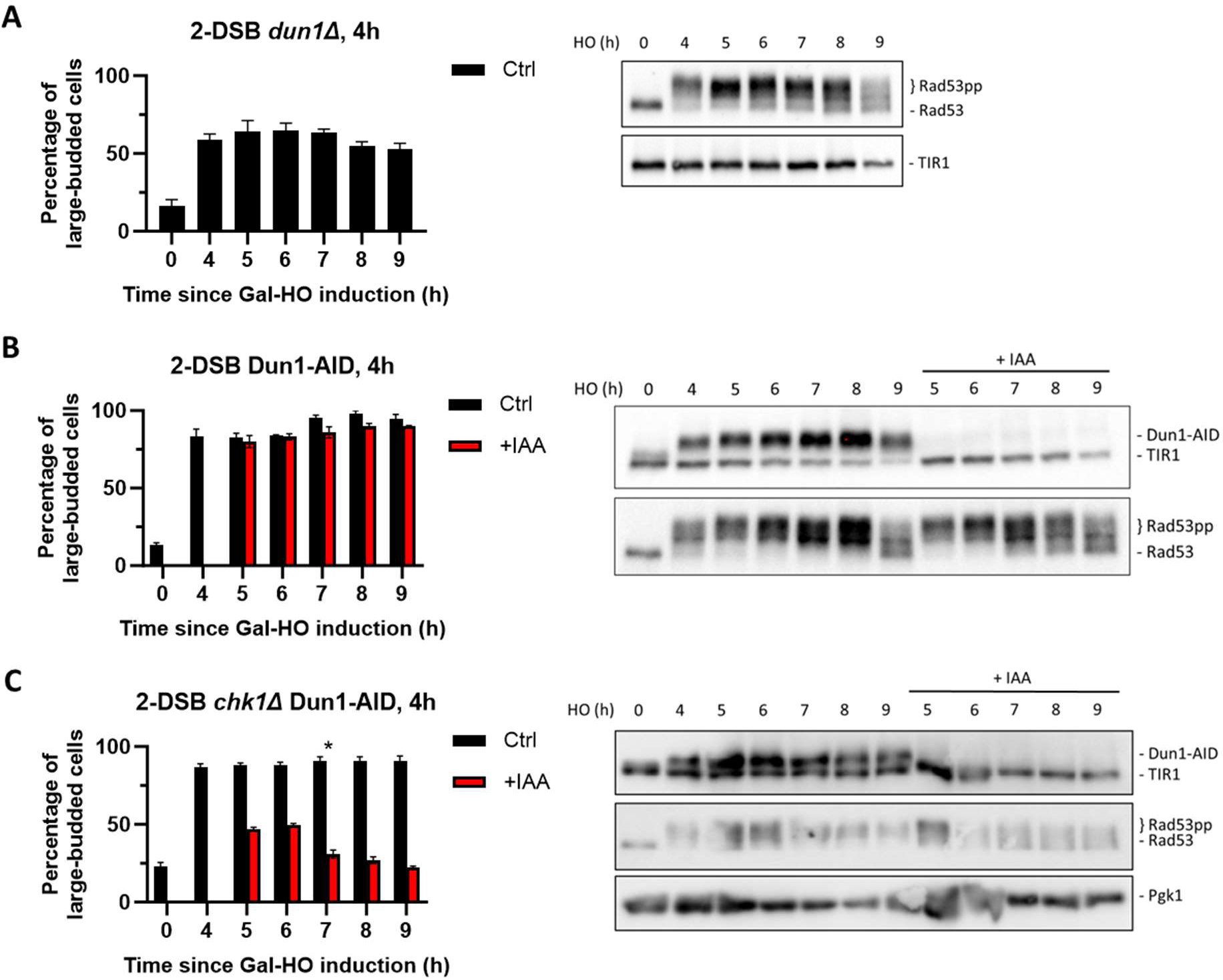
Dun1 is not required for checkpoint maintenance. (A) Adaptation assay of 50 G_1_ cells on a YEP-Gal plate with 2-DSB *dun1Δ*. G_2_/M arrest was determined based on cell morphology as shown in Figure 1A. Data is shown from 3 trials with standard error of the mean (SEM). Western blot probed with α-Rad53 and α-Myc for TIR1-Myc as a loading control. (B) Percent G_2_/M arrested cells for 2-DSB *DUN1-AID* after HO induction. Data are shown from 3 trials with standard error of the mean (SEM). Cultures were split 4 h after DSB induction; with auxin (1 mM) (+IAA). Western blot probed with α-Rad53 and α-Myc. α-Rad53 shows both an unphosphorylated protein and multiple phosphorylated species. α-Myc shows Dun1-AID degradation and TIR1-Myc as a loading control. (C) Same as (B) for 2-DSB *chk1Δ DUN1-AID*. The asterisk marks when the percentage of large-budded cells returned to pre-damage levels.

### Ddc2 and Rad53’s Role in Maintaining Arrest Become Dispensable in Extended G_2_/M Arrest

Previously, we reported that Ddc2 protein abundance initially increases over time as after the induction of a single DNA break, which is followed by near-complete depletion of Ddc2 around the time that cells adapt (Memisoglu et al. 2019). Given that Ddc2 overexpression leads to permanent cell cycle arrest following DNA damage (Clerici et al. 2001), we concluded that Ddc2 abundance is intimately tied to the duration of the arrest. Here, we asked whether the presence of 2 DSBs instead of a single DSB would lead to an increase in Ddc2 protein abundance and therefore, the permanent cell cycle arrest. We examined the levels of Ddc2 in both 1 and 2-DSB strains following DNA damage but did not detect a difference in the changes of abundance of Ddc2 protein (**Figure S5A-C**), even though cells adapt to 1 DSBs and remained terminally arrested after 2 DSBs (**Figure 1A-1D**).

If Ddc2 activity is essential to maintain the cell cycle arrest in the 2-DSB strain, then degradation of Ddc2-AID around the time wild-type cells adapt to a single DNA break should interrupt the permanent cell cycle arrest and trigger cell cycle re-entry. We find that complete depletion of Ddc2-AID 15 h after the induction of 2 DSBs leads to diminished Rad53 hyperphosphorylation (**Figure 5A**). However, surprisingly, Ddc2-AID degradation did not alter the percentage of G_2_/M arrested cells even 9 h after Ddc2 depletion (**Figure 5A).** Furthermore, despite the diminished Rad53 phosphorylation, Ddc2-AID cells mostly remained arrested in G_2_/M after the depletion of Ddc2-AID at 15 h as illustrated by DAPI staining (**Figure 5B**), in contrast to Ddc2-AID depletion soon after induction of 2 DSBs which causes cells to rapidly resume mitosis (**Figure 2A**). These results hint that the maintenance of the permanent cell cycle arrest in response to 2 DSBs at later stages could be independent of the DDC signaling.

**Figure 5.**
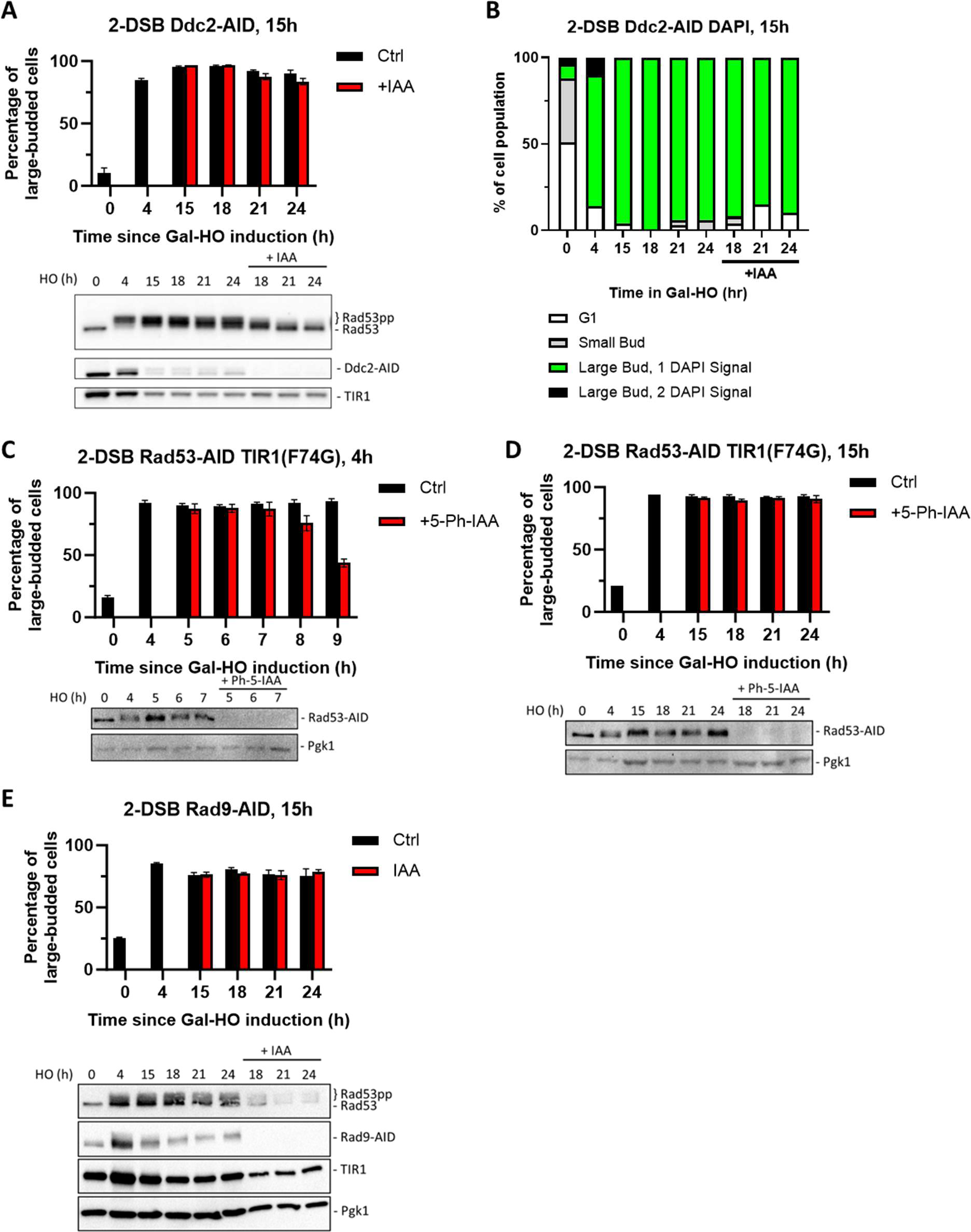
Ddc2 and Rad53 are dispensable for >24 h checkpoint arrest. (A) Percent G_2_/M arrested cells for 2-DSB *DDC2-AID* after HO induction. Data is shown from 3 trials with standard error of the mean (SEM). Western blot probed with α-Rad53 and α-Myc. α-Rad53 shows both an unphosphorylated protein and multiple phosphorylated species. α-Myc shows Ddc2-AID degradation and TIR1-Myc as a loading control. (B) Profile of DAPI stained cells in a 2-DSB *DDC2-AID* strain after HO induction. Cells were categorized based on cell morphology and number of DAPI signals. (C) Percent G_2_/M arrested cells for 2-DSB *RAD53-AID TIR1(F74G)* after HO induction. 5-Ph-IAA was added 4 h after HO induction. Data is shown from 3 trials with standard error of the mean (SEM). Western blot probed with α-Rad53, α-Myc, and α-Pgk1. α-Rad53 shows both an unphosphorylated protein and multiple phosphorylated species. α-Myc shows Rad53-AID degradation. α-Pgk1 probed as a loading control. (D) Same as (C) where 5-Ph-IAA was added 15 h after HO induction. (E) Percentage G_2_/M arrested cells for 2-DSB *RAD9-AID* plus pRad9-AID after HO induction. Data shown from 3 trials with standard error of the mean (SEM). Western blot probed with α-Rad53 and α-Myc. α-Rad53 shows both an unphosphorylated protein and multiple phosphorylated species. α-Myc shows Rad9-AID degradation and TIR1-Myc as a loading control. α-Pgk1 probed as a loading control.

We then asked whether other DDC factors such as Rad9, Rad24 and Rad53 are dispensable for the prolonged arrest following the induction of 2 DSBs. However as noted above, AID-tagged DDC activation proteins were unable to maintain this prolonged arrest in a 2-DSB strain even in the absence of IAA, which precludes their use in this analysis. To be able to study the contribution of these DDC factors in prolonged cell cycle arrest, we turned to the AID version 2 (AID2) system (Yesbolatova et al. 2020). To this end, we integrated a TIR1-F74G point mutation, and used 5-phenyl-IAA (5-Ph-IAA) instead of IAA to lower the basal degradation of AID-tagged proteins. Switching to the AID2 system did not fully restore function to *RAD9-AID2* and *RAD24-AID2* strains, as they mostly adapted after 24 h after the exposure to DNA damage, but 87% of *RAD53-AID2* cells remained in G_2_/M arrest at 24 h (**Figure S1C**). Using Rad53-AID2, we then asked whether Rad53 is required for extended G_2_/M arrest in response to 2 DSBs. Rapid depletion of Rad53-AID2 with 5-Ph-IAA 4 h after DSB induction led cells to escape G_2_/M arrest, but as with the *RAD53-AID* strain, we detected a 4 h delay in cell cycle re-entry, confirming our previous results **(Figure 5C and Figure 2D**). However, degradation of Rad53-AID2 15 h after DSB induction did not prompt cell cycle re-entry even 9 h after complete depletion of Rad53 (**Figure 5D, S. Table 1**), akin to what we observe following the depletion of Ddc2-AID (**Figure 5A-5B**).

Because TIR1-mediated degradation of Rad9-AID even without auxin caused most cells to adapt 24 h after inducing DNA damage, we asked whether overexpression of *RAD9-AID* could overcome this effect. We added a *TRP1* centromere-containing plasmid copy of *RAD9-AID* with its endogenous promoter (pRAD9-AID) to our 2-DSB *RAD9-AID* strain. Degradation of Rad9-AID by IAA 15 h after DSB induction did not trigger release of cells from G_2_/M arrest (**Figure 5E, S. Table 1**). However, unlike degradation of Ddc2-AID or Rad53-AID2 in this same situation, Rad53 remained hyperphosphorylated up to 9 h after adding IAA (**Figure 5E**). Therefore, while the DDC proteins Ddc2, Rad9, and Rad53 are required for the maintenance of checkpoint arrest at early stages, surprisingly, they are dispensable for prolonged arrest following induction of 2 DSBs. These results suggest that prolonged cell cycle arrest is maintained by signaling proteins other than the Mec1 branch of the DDC.

### Spindle Assembly Checkpoint Proteins Mad1 and Mad2 are Required for Prolonged Arrest

In addition to the DDC, the spindle assembly checkpoint (SAC) maintains genomic integrity by halting mitosis at the metaphase/anaphase transition in response to unattached kinetochores, to ensure accurate chromosome segregation (reviewed by (McAinsh and Kops 2023)). We have previously shown that inactivation of the SAC by a *MAD1, MAD2* or *MAD3* deletion shortened the duration of the cell cycle arrest induced by a single DSB (Dotiwala et al. 2010). To test whether SAC is involved in enforcing and sustaining permanent cell cycle arrest in response to 2 DSBs, we deleted *MAD2* in the 2-DSB strain. Adaptation assay results illustrate that nearly all *mad2Δ* cells arrested at 4 h but began to adapt between 12-15 h after DNA damage (**Figure S6A**). To explore whether deletion of *MAD2* can antagonize the permanent cell cycle arrest due to hyperactive DDC signaling, we overexpressed Ddc2 in *mad2Δ* 2-DSB cells and assayed mitotic progression. We find that both in 1-DSB and 2-DSB strains, *MAD2* deletion leads to cell cycle re-entry even when Ddc2 is overexpressed (**Figure S6B**). Based on these data, we concluded that mitotic inhibition is enforced by the SAC proteins as DDC factors become dispensable 12 to 15 h after the induction of damage.

If SAC proteins are only required at later stages of cell cycle arrest when DDC proteins are dispensable, then the depletion of SAC protein Mad2 or its binding partner Mad1 soon after the induction of DNA damage should not affect DDC or cell cycle arrest up to 12 to 15 hours. In agreement with this idea, after Mad1-AID or Mad2-AID depletion at 4 h, cells remained arrested up to 15 h following DSB induction, with persistent Rad53 hyperphosphorylation (**Figure S7A-C**). Strikingly, these cells eventually re-entered the cell cycle by 24 h (20 h after depletion of Mad1 and Mad2). It is notable that cells resumed cell cycle progression despite persistent Rad53 hyperphosphorylation. This result reinforces our conclusion that Mad1 and Mad2 are not required for the activation and initial maintenance of arrest but are essential for prolonged cell cycle arrest in response to 2 DSBs. These data also suggest that in the absence of Mad1 or Mad2, cells become insensitive to the arrest normally imposed by DDC.

To monitor the effect of SAC proteins Mad1 and Mad2 at late stages of prolonged cell cycle arrest when DDC signaling becomes dispensable, we depleted Mad1-AID or Mad2-AID 15 h after DSB induction. Following the depletion of Mad1-AID and Mad2-AID, we observed an immediate reduction in the percentage of G_2_/M-arrested cells (**Figure 6A-C**). DAPI staining confirmed that Mad1-AID or Mad2-AID deplete cells re-entered the cell cycle, evident from an increase in the G1 cell population as well as an increase in percentage of large-budded cells with 2 DAPI foci (**Figure 6B, D**). Similar to what we observe following Mad1-AID and Mad2-AID depletion 4 h after damage, this cell cycle re-entry occurred despite persistent Rad53 hyperphosphorylation (**Figure 6A, C**).

**Figure 6.**
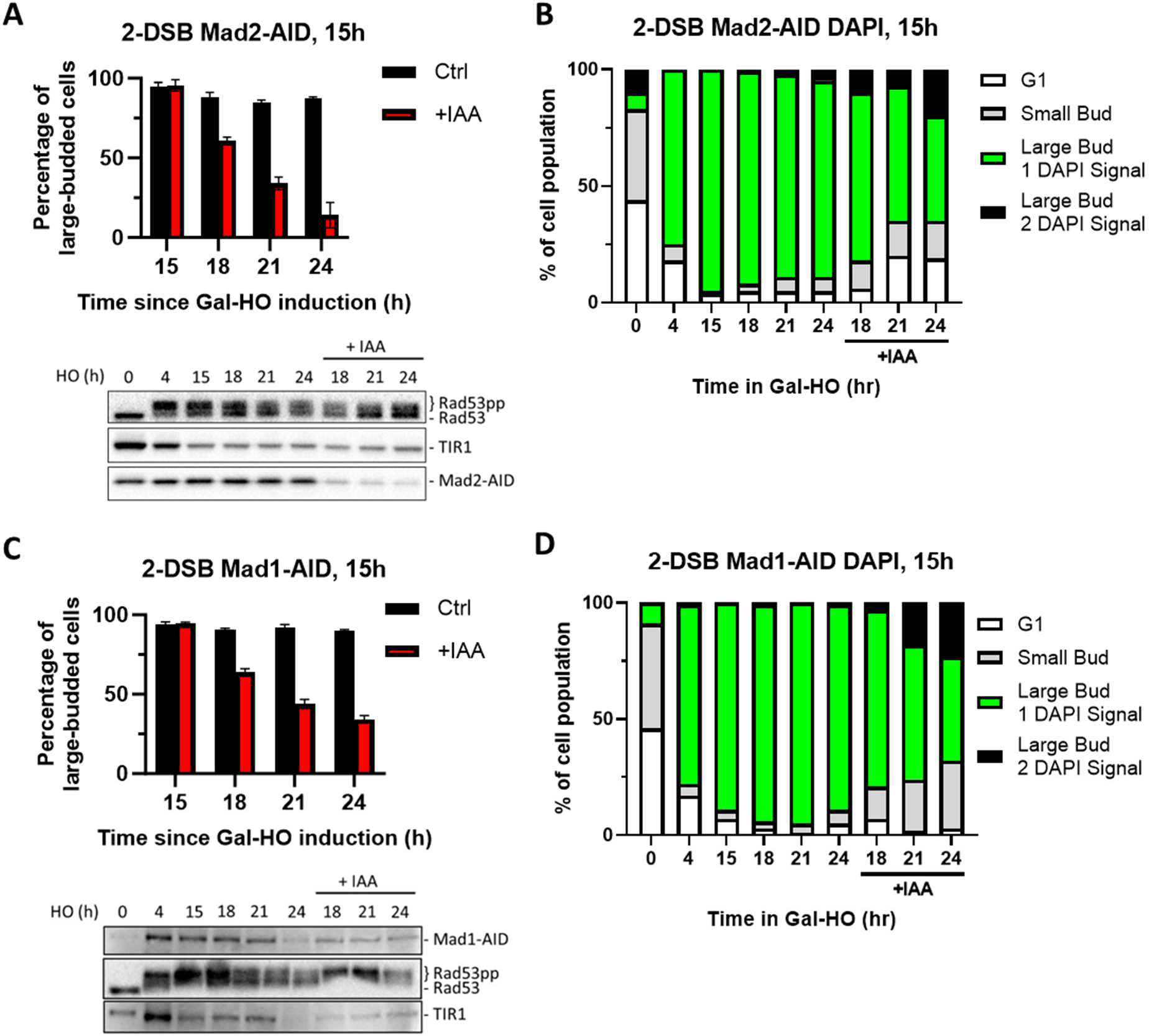
Degradation of Mad2 or Mad1 at 15 h releases cells from checkpoint arrest. (A) Percent G_2_/M arrested cells for 2-DSB *MAD2-AID* after HO induction. Data is shown from 3 trials with standard error of the mean (SEM). Western blot probed with α-Rad53 and α-Myc. α-Rad53 shows both an unphosphorylated protein and multiple phosphorylated species. α-Myc shows Mad2-AID degradation and TIR1-Myc as a loading control. (B) Profile of DAPI stained cells in a 2-DSB *MAD2-AID* strain after HO induction. Liquid cultures were split 15 h after HO induction and treated with either IAA or ethanol. Cells were scored based on cell morphology and number of DAPI signals. (C) Same as (A) for 2-DSB *MAD1-AID*. (D) Same as (B) for 2-DSB *MAD1-AID*.

To show that the late stages of the permanent cell cycle arrest in response to 2 DSBs is independent of DDC and dependent on SAC, we inactivated DDC by depleting Ddc2-AID together with Mad2-AID. Although the simultaneous depletion of Ddc2-AID and Mad2-AID led to Rad53 dephosphorylation as detected by western blotting, we found no statistically significant difference in the percentage of cells escaping G_2_/M arrest in response to *DDC2-AID MAD2-AID* double depletion strain compared to Mad2-AID alone (**Figure S8A-B, and S. Table 1**). Collectively, our findings indicate that DDC initiates and sustains the cell cycle arrest approximately for 15 h following DNA damage, but after that DDC becomes dispensable and the permanent arrest is sustained by SAC.

### Mitotic Exit Network Proteins Bfa1 and Bub2 have Different Roles in the DDR

To investigate the possible contribution of the mitotic exit network (MEN) to the maintenance of the extended cell cycle arrest in response to 2 DSBs, we appended AID tags to upstream MEN proteins Bub2 and Bfa1. Degradation of Bub2-AID both 4 h and 15 h after induction of 2 DSBs suppressed G_2_/M arrest (**Figure 7A, S9A-B**), akin to what we observe following Mad1-AID or Mad2-AID depletion (**Figure 6A-B**). In contrast to Bub2-AID, we see that Bfa1-AID degradation did not trigger a significant release from G_2_/M arrest and did not alter the phosphorylation of Rad53 (**Figure 7B, S9C-E**). Analysis of cell cycle distribution with DAPI staining showed that neither the inactivation of Bub2 nor Bfa1 led to the accumulation of a significant number of cells with two separate DAPI-staining nuclei, which is indicative of mitotic exit defects (**Figure 7B, D and S9B, D, F**). These results imply that although Bub2 and Bfa1 have interdependent functions for MEN signaling, they carry out independent roles in response to DNA damage.

**Figure 7.**
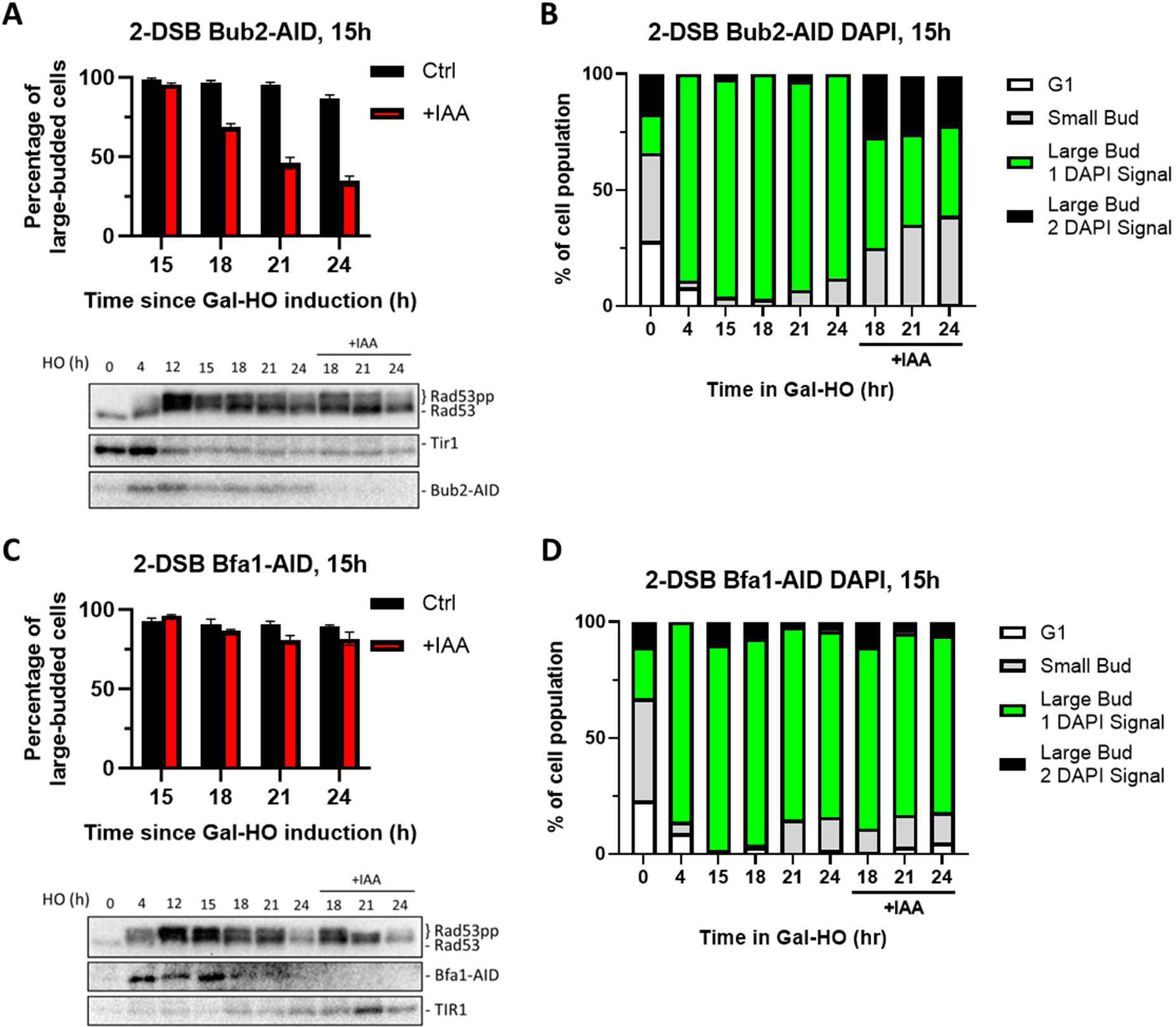
Degradation of Bub2 but not Bfa1 at 15 h releases cells from checkpoint arrest. (A) Percent G_2_/M arrested cells for 2-DSB *BUB2-AID* after HO induction. Data is shown from 3 trials with standard error of the mean (SEM). Western blot probed with α-Rad53 and α-Myc. α-Rad53 shows both an unphosphorylated protein and multiple phosphorylated species. α-Myc shows Bub2-AID degradation and TIR1-Myc as a loading control. (B) Profile of DAPI stained cells in a 2-DSB *BUB2-AID* strain after HO induction. Liquid cultures were split 15 h after HO induction and treated with either IAA or ethanol. Cells were scored based on cell morphology and number of DAPI signals. (C) Same as (A) for 2-DSB *BFA1-AID*. (D) Same as (B) for 2-DSB *BFA1-AID*.

### The Location of the Second DSB Site Relative to the Centromere Affects Prolonged Arrest

In contrast to the permanent cell cycle arrest we observe in response to 2 DSBs, a recent study using two HO-mediated persistent DSBs showed that cells in fact adapt (Sadeghi et al. 2022). One difference between these two 2-DSB systems is the relative position of the DSBs, which might affect how SAC components become engaged, and thus might determine the extent of mitotic arrest. Supporting this, we previously showed that deleting *CEN3* in a strain with a DSB at *MAT* on chromosome III eliminated the Mad2-dependent delay in adaptation, but deleting *CEN3* when the DSB was on chromosome VI had no effect (Dotiwala et al. 2010). In our adaptation-defective 2-DSB strain, the DSBs are located at *MAT* (86 kb from *CEN3*) and near *FAB1* (42 kb from *CEN6*). Sadeghi et al. employed 2 strains, both of which escape prolonged G_2_/M arrest, with at least one DSB site far from its centromere; at *URA3* (36 kb from its centromere) and *ADH1* (170 kb) or at *MIC2* (32 kb) and *DLD2* (316 kb).

To investigate whether the distance between the second DSB site and the centromere would affect whether cells will remain permanently arrested, we created several strains which contain a second cut site at various distances from the centromere, in addition to the cut site at *MAT* on chromosome III (**Figure S10**). Strain YSL53, which has a second DSB at chromosome V, 86 kb away from *CEN5* (Lee et al. 1998), and strain DW417, which has a second DSB 52 kb away from *CEN6* (Lee et al. 2014) mostly remained in G_2_/M arrest 24 h after the induction of DNA breaks. However, in GEM188, which had a second cut site 230 kb away from the *CEN2*, only 37% of cells remained in G_2_/M arrest by 24 h. Thus, the increased distance of the second DSB site to the centromere in GEM188 appears to have led to a less robust triggering of the SAC compared with YSL53 and DW417.

Our previous data have suggested that the involvement of the SAC in prolonging DSB-induced arrest was dependent on centromere sequences on the broken chromosome and involved post-translational modification of chromatin by the Mec1- and Tel1-dependent phosphorylation of the histone H2A (Dotiwala et al. 2010). In budding yeast, histone H2B is also targeted by DDC kinases upon DNA damage (Lee et al. 2014). To test whether the presence of these chromatin modifications around centromeres would be sufficient to elicit a SAC response, we examined cell cycle progression in strains in which both histone H2A and/or histone H2B genes were mutated to their putative phosphomimetic forms (H2A-S129E and H2B-T129E). We note that although histone H2A-S129E is recognized by an antibody specific for the phosphorylation of histone H2A-S129 (Eapen et al. 2012), the mutation to S129E may not be fully phosphomimetic. As shown in Figure S11, there was no effect on the growth rate of either the single or the double mutants, suggesting that cells did not experience a SAC-dependent delay in entering mitosis because of these modifications.

## Discussion

Here, we studied how cells maintain cell cycle arrest following DNA damage, by using a yeast strain that permanently halts cell cycle with two persistent DNA breaks. We find that most of the DDC signaling proteins must remain active to maintain G_2_/M arrest, highlighting the importance of continuous checkpoint signaling in preventing premature mitotic entry and therefore, genome instability (**Figure 8**).

**Figure 8.**
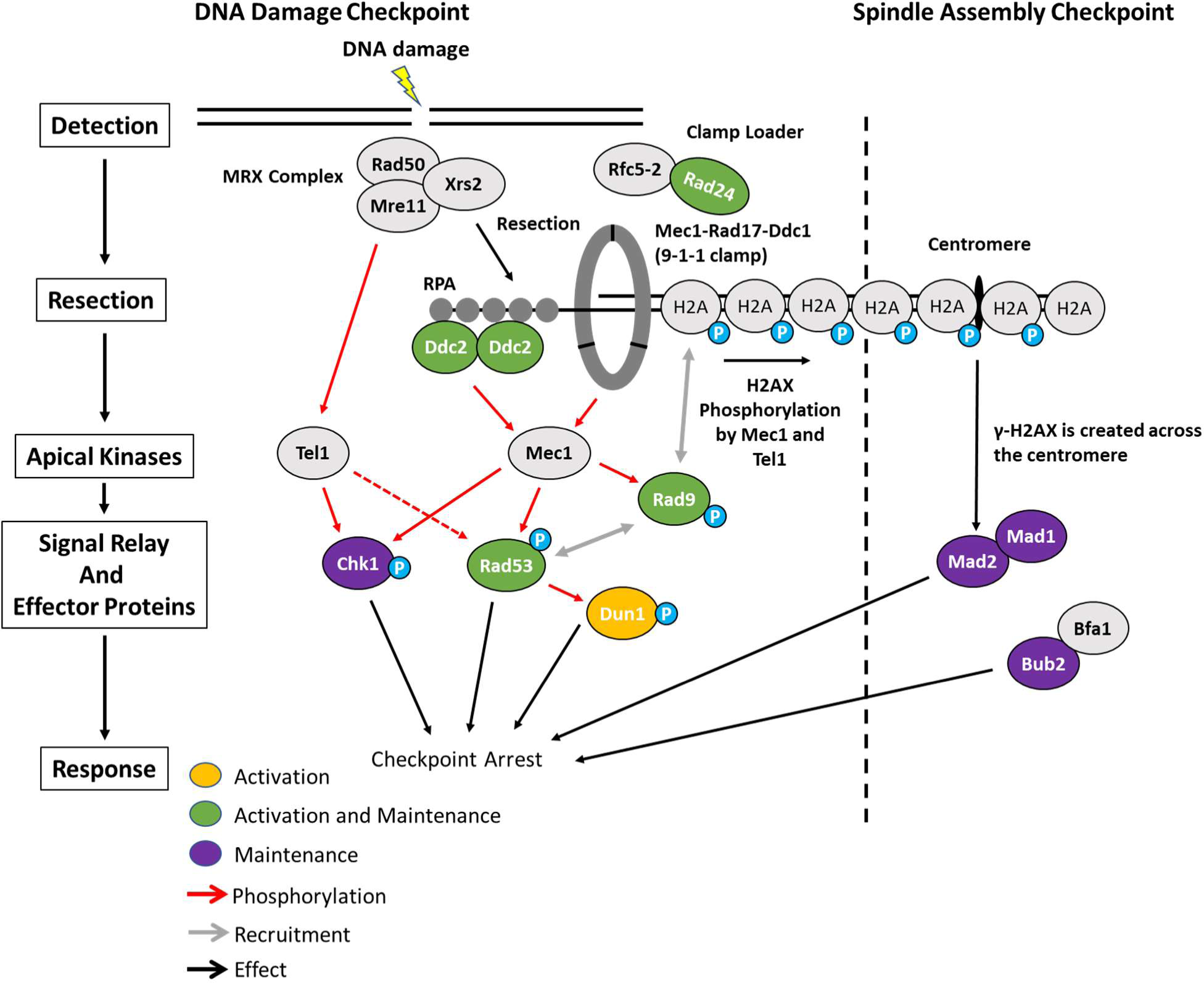
Activation and maintenance of checkpoint arrest in response to a DSB. The Mre11-Rad50-Xrs2 (MRX) complex is one of the first complexes recruited to DSBs and initiates the resection of dsDNA to ssDNA. ssDNA is then coated with RPA which recruits Ddc2. Mec1 is the primary kinase responsible for checkpoint arrest in budding yeast and is activated by Ddc2 and Ddc1 from the 9-1-1 clamp. Proteins in green (Ddc2, Rad9, Rad24, and Rad53) were required for the activation and maintenance of checkpoint arrest. While Chk1 was not required for establishment of G_2_/M arrest, it contributed to the maintenance of arrest. In contrast, Dun1 was required for checkpoint activation but was dispensable 4 h after DSB induction. Prolonged arrest >24 h in a 2-DSB strain was dependent on the SAC proteins Mad2, Mad1, and Bub2 as well as the distance between the 2^nd^ HO-cut site and the centromere.

Phosphorylation of Rad53 by Mec1 is tightly linked to cell cycle arrest following DNA damage (Pellicioli et al. 2001). While Rad53 is also shown to be targeted by Tel1, deleting *TEL1* did not affect the prolonged checkpoint arrest in a 2-DSB strain. This finding supports the previous reports showing that cell cycle arrest in response to enzymatic DNA breaks is largely orchestrated by Mec1, with a minor contribution from Tel1 (Pellicioli et al. 2001; Vaze et al. 2002). Here, we add that Mec1 inactivation via Ddc2 depletion, 4 h after DSB induction, results in rapid resumption of the cell cycle, most likely through Ptc2, Ptc3, Pph3 and Glc7-dependent dephosphorylation and inactivation of the DDC signaling protein Rad53 (Bazzi et al. 2010; Leroy et al. 2003; O’Neill et al. 2007). However, the regulation and consequences of Rad53 phosphorylation are apparently more complex, given that cells can re-enter mitosis in the presence of persistent Rad53 phosphorylation when SAC proteins Mad1 and Mad2 are depleted (see below).

Mec1 activation in the S and G_2_ cell cycle phases is achieved by at least two converging mechanisms; first, through Ddc2 binding to RPA-coated ssDNA created the 5’ to 3’ resection of the DSB ends, and second, through binding of the 9-1-1 clamp subunit Ddc1 (Dubrana et al. 2007; Navadgi-Patil and Burgers 2009; Zou and Elledge 2003; Melo, Cohen, and Toczyski 2001). Before Ddc1 can activate Mec1, the 9-1-1 checkpoint clamp must be loaded by the clamp loader, which consists of Rad24-Rfc2-5 (Melo, Cohen, and Toczyski 2001). DDC activation largely depends on a functional clamp loader, as cells lacking *RAD24* proceed directly into mitosis in response to a single DSB, with only a brief delay (Aylon and Kupiec 2003). If the clamp loader acts just once to load the 9-1-1 clamp and the clamp then slides away from the DSB as DNA is resected, then removal of Rad24 after the checkpoint had been robustly activated should not perturb arrest. However, we find that Rad24 depletion leads to rapid cell cycle resumption in response to two DSBs, suggesting that multiple 9-1-1 clamp loading events are required to sustain extended cell cycle arrest. We posit that 9-1-1 clamp may require continuous reloading, as the 5’ end is being continuously resected by Exo1 and Sgs1-Rmi1-Top3-Dna2 exonucleases (Zhu et al. 2008).

The adaptor kinase Rad9, downstream of Mec1 and Tel1, is responsible for scaffolding and activating the effector DDC kinases Rad53 and Chk1 (Emili 1998; Schwartz et al. 2002; Sweeney et al. 2005). Here, we show that conditional depletion of Rad9 shortly after the induction of 2 DSBs prompts mitotic re-entry and terminates the DDC, evident from rapid Rad53 dephosphorylation. Thus, Rad9 is required for continued maintenance of Rad53 phosphorylation.

Depletion of Rad53-AID 4 h after inducing 2 DSBs triggers release of cells from G_2_/M arrest, albeit resumption of mitosis is delayed by 4 h compared to resumption of mitosis seen when upstream DDC factors Ddc2, Rad9, or Rad24 are depleted. We discovered that this residual delay in mitotic re-entry is dependent on Chk1 signaling. We postulate that Chk1 may delay cell cycle re-entry in the absence of Rad53 by phosphorylating and stabilizing Pds1 (Agarwal et al. 2003).

Agreeing with previously published work (Yam et al. 2020), we find that deleting *DUN1* resulted in a less robust DDC activation and a shortened G_2_/M arrest. However, unlike the depletion of other DDC proteins examined in this study, depletion of Dun1-AID 4 h after DSB induction was not sufficient to promote cell cycle re-entry. It is possible that the transcripts upregulated by Dun1 after DNA damage (Zhao and Rothstein 2002; Zhou and Elledge 1993) are stable for several hours and are sufficient to sustain the arrest even in the absence of Dun1. We also find that cell cycle arrest upon depletion of Dun1 is Chk1-dependent. These results are in consistent with Dun1’s role in stabilizing Pds1 through a Chk1-independent mechanism (Yam et al. 2020).

Here, we show that prolonged cell cycle arrest following induction of 2 DSBs becomes independent of DDC proteins Ddc2, Rad9, and Rad53, but dependent on SAC proteins Mad1 and Mad2. Depletion of Mad1 or Mad2 4 h after DSB induction did not immediately result in cell cycle re-entry, but around 15 h cells began to resume cell cycle. The timing of cell cycle re-entry for Mad1/2-depleted cells following the induction of 2 DSB is about the same as the timing of cell cycle re-entry in response to a single DNA break in wild-type cells. Even when DDC signaling is artificially upregulated via Ddc2 overexpression, cells re-enter mitosis if Mad2 is depleted. Surprisingly, in the absence of Mad1 or Mad2 cells escaped arrest despite persistent Rad53 hyperphosphorylation, suggesting that cells become “deaf” to the DDC signal once SAC takes over.

Previous work indicates that SAC proteins contribute to the DNA damage response (Dotiwala et al. 2010; Garber and Rine 2002; Kim and Burke 2008). Work from our lab has suggested that γ-H2AX spreading from a DSB to the centromere of the same chromosome might impair kinetochore attachment and thus trigger a SAC response (Dotiwala et al. 2010). SAC signaling is not sufficient to elicit a permanent cell cycle arrest in response to a single DSB; however, we find that inducing 2 DSBs, each within 100 kb of its centromere, elicits a SAC-dependent permanent arrest. As strength of the SAC has a direct relation to the number of unattached kinetochores (Dick and Gerlich 2013), the addition of a second DSB on another chromosome might trigger a stronger SAC response and result in a permanent cell cycle arrest.

The fact that not all combinations of two DSBs produce permanent arrest (Sadeghi et al. 2022) can be explained by the idea that the distance between the DSB site and its corresponding centromere is an important determinant of the extent of cell cycle arrest. We previously observed that a strain with a single DSB 200 kb away from the centromere had shorter cell cycle arrest compared to a strain with a DSB 86 kb away from the centromere (Dotiwala et al. 2010). Here we provide further evidence suggesting that the distances between DSBs and their corresponding centromeres determine whether SAC will be fully activated to prolong the arrest. Our previous results showed that when *MAD2* was deleted, the length of cell cycle arrest was the same in strains with a single DSB, irrespective of the DSB’s distance to its centromere (Dotiwala et al. 2010). We suggest that the strains used by (Sadeghi et al. 2022) do not remain permanently arrested because one of the two DSBs in their strains is sufficiently far from its centromere to fully trigger SAC.

In addition to blocking the metaphase to anaphase transition, components of the SAC also block mitotic exit (reviewed by (Matellan and Monje-Casas 2020)). We examined the role of Bub2/Bfa1 heterodimer, the most upstream components of the MEN pathway (Matellan and Monje-Casas 2020). Much like Mad1 and Mad2, we found that neither Bub2 nor Bfa1 was required for the establishment of cell cycle arrest in response to DNA damage; but our study revealed a surprising result: Bub2, but not its partner Bfa1, is essential to prolong cell cycle arrest, indicating that Bub2 has a Bfa1-independent role.

By using conditional depletion of various proteins that contribute to cell cycle arrest, we show that the establishment, maintenance, and inactivation stages of DNA damage-provoked cell cycle arrest involve different sets of factors following DNA damage. After the DNA damage checkpoint is established, its maintenance proves to be divided into two distinct phases. Arrest up to about 15 h requires the constant presence of most of the identified DDC proteins, including Ddc2, Rad9, Rad24, and Rad53, with Dun1 playing an important but nonessential role. Although Chk1 was not required either to establish or initially to maintain cell cycle arrest, its absence shortened arrest, most notably when Rad53 was depleted. Surprisingly, neither Ddc2, Rad9, nor Rad53 (and we suggest likely Rad24) are necessary for the prolongation of cell cycle arrest lasting longer than 15-24 h. Instead, this prolonged arrest is enforced by SAC proteins Mad1, Mad2, and Bub2.

## Methods and Materials

### Yeast Strain and Plasmid Construction

All AID-tagged mutant strains were derived from a modified version of strains JKM179 (**S. Table 2**). To create the strain with two HO cleavage sites (DW417), an HO cut site, designated HOcse6, with an adjacent HPH marker was integrated into chromosome VI, 52 kb from the centromere. To create auxin-inducible degron strains, we first integrated osTIR1 at *URA3* after digesting plasmid pNHK53 (Nishimura et al. 2009) with *Stu*I. To integrate osTIR1-F74G at *URA3*, the plasmid pMK420 (Yesbolatova et al. 2020) was digested with *Stu*I. For degron-tagging of DDC proteins, AID-9xMyc (AID) PCR products were generated with mixed oligos with homology to the C-terminal end of the corresponding open reading frames by using plasmids pKan–AID–9xMyc (pJH2892) or pNat–AID–9xMyc (pJH2899) as templates (Morawska and Ulrich 2013). Deletion of ORFs and insertion of AID tags were introduced with the one-step PCR homology cassette amplification and the standard yeast transformation method (Wach et al. 1994). Cas9 editing was done by inserting a gRNA into plasmid bRA90 (Anand et al. 2017) and co-transformed into our strain of interest with a donor sequence. Transformants were verified by PCR, western blotting, and sequencing. To create strain GEM188 with 2-DSBs, we inserted a second HO-cut site into JKM179 at *LYS2* locus by CRISPR/Cas9 (Anand et al Protocol Exchange, 2017) using a synthetic DNA template with 117 bp consensus HO recognition site. Non-phosphorylatable and phosphomimetic mutants of H2A were generated in a JKM179 background using CRISPR/Cas9 to target HTA1 and HTA2 genes at serine 129 and 80nt templates to mutate serine to either alanine (non-phosphorylatable) or glutamic acid (phosphomimetic). H2B mutants were generated in a JKM179 background using CRISPR/Cas9 to target *HTB1* and *HTB2* genes at threonine 129 with 80nt repair templates to mutate threonine to either alanine or glutamic acid.

The *CEN/ARS* plasmid pFZ052-*pRAD9-AID-*9Myc::Trp1* (pRAD9-AID) was obtained by digesting the plasmid pFL36.1 (Lazzaro et al. 2008) with *SmaI* and *AscI* to excise the *3-HA* tag on the C-terminal end of Rad9. A *9xMyc-AID* PCR product generated from the plasmid pJH2892 (pKan-9xMyc-AID) was cut with *AscI* and sticky/blunt end cloned into the *SmaI-AscI*-digested pFL36.1 to add the *9xMyc-AID* tag to the C-terminal end of Rad9. pRad9-AID was retained by growing cells in a Trp-media with 2% raffinose. Primers used for strain and plasmid creation are listed in **S. Table 3**. Plasmids are listed in **S. Table 4**.

### Culturing Conditions, HO expression, and Auxin Treatment

Strains containing degron fusions and galactose-inducible HO were cultured using standard procedures. Briefly, a single colony grown on a YEPD plate (1 % yeast extract, 2 % peptone, 2 % dextrose, 2.5 % agar) was inoculated in 5 ml YEP-lactate (YEP containing 3 % lactic acid) and was grown for ∼15 hours at 30° C with agitation. Next day, the overnight culture was used to inoculate a 500-100 ml of YEP-lactate culture such that the cell density reached an OD_600_ of 0.5 the following day. After harvesting 15 ml liquid culture before treatment, HO expression was induced by galactose treatment with a 2% final concentration. Then, cultures were split either at 4 h or 15 h following induction with galactose. The split cultures were treated either with IAA or 5-Ph-IAA or an equivalent volume of 200 proof ethanol. Indole-3-acetic acid (IAA, Sigma-Aldrich, I3750) was resuspended in ethanol for a 500 mM stock and used at a 1 mM final concentration both for liquid media and for agar plates. 5-Ph-IAA was dissolved in ethanol for a 1 mM stock and used at a final concentration of 1µM. 15 ml of liquid culture were harvested at various time points and prepared for microscopy or western blot analysis as described below.

### Culturing Conditions, HO expression, and Auxin Treatment

To measure growth rate, strains were grown in 5 ml of YEPD with 2% dextrose at 30 °C with an initial OD_600_ of 0.1. The OD was measured 3, 5, 7, and 10 h after the initial OD measurement using a Thermo Scientific NanoDrop 2000c Spectrophotometer. To measure the OD, 50µl of culture was added to 950µl of fresh YPD in a cuvette at a dilution of 1:20.

### TCA Protein Extraction

Protein extracts were prepared for western blot analysis by TCA extraction protocol as previously explained (Miller-Fleming et al. 2014). Briefly, 15 ml of liquid culture was spun down and the media was discarded. Harvested cells were incubated on ice in 1.5 ml microcentrifuge tubes with 20% TCA for 20 minutes. Cells were washed with acetone and the pellet was air dried. 200 µl of MURBs buffer (50 mM sodium phosphate, 25 mM MES, 3M urea, 0.5% 2-mercaptoethanol, 1 mM sodium azide and 1% SDS) was added to each sample with acid washed glass beads. Cells were lysed by mechanical shearing with glass beads for 2 min. Crude cell lysates were harvested by poking a hole in the bottom of the 1.5 ml microcentrifuge tube and spinning the tubes on a 15 ml conical tube. Samples were boiled at 95 ^0^C for 10 min prior to loading on SDS-PAGE.

### Western Blotting

8 – 20 µl of denatured protein samples prepared by TCA extraction were loaded onto a 10 % or 8 % SDS-PAGE gels. Proteins were separated by applying constant voltage at 90 V until the 37 kDa protein standard band reached the bottom of the gel. Gels were transferred to an Immun-Blot PVDF membrane (BioRad) using a wet transfer apparatus set to 100 V constant voltage for 1 h at 4 °C. Membranes were then blocked with OneBlock blocking buffer (Genesee Scientific Cat #: 20-313) for 1 h at room temperature, After three 10 minute washes with 1x TBS-T, blots were incubated with either anti-Myc [9E11] (Abcam, ab56) to detect TIR1 and AID fusions, anti-Rad53 [EL7.E1] (Abcam, ab166859), anti-Pgk1 (Abcam, ab30359), or anti-Rad9 (Usui, Foster, and Petrini 2009) for 1 h at room temperature or at 4 ^O^C overnight. Blots were washed 3 times with 1x TBS-T and incubated with anti-mouse HRP (GE Healthcare Cat # NXA931) or anti-rabbit HRP secondary antibody (Sigma-Aldrich, Cat # A6154) for 1 h at room temperature. After washing the membranes 3 times with 1x TBS-T, Amersham ECL Prime Western Blotting Detection Reagent was added to fully coat the blots and left to incubate for 5 min at room temperature with gentle agitation. Blots were imaged using a BioRad ChemiDoc^TM^ XR+ imager and prepared for publication using ImageLab 6.1 software (BioRad) and Adobe Photoshop CC 2017. Reagents used are listed in **S. Table 5**.

### Microscopy, DAPI staining, and Cell Morphology Determination

Aliquots from YEP-Lac cultures were taken either 4 h or 15 h after adding galactose, diluted 20-fold in sterile water and plated on a YEP-Agar with 2 % galactose with or without 1 mM IAA or 1 µM 5-Ph-IAA. Cells were counted on a light microscope with a 10x objective, examined and binned into three categories: unbudded, small buds, and G_2_/M arrested cells with large buds. For each time-point, >250 cells were analyzed. For DAPI staining, 450 µl of culture was added to 50 µl of 37% formaldehyde and incubated in the chemical hood at room temperature for 20 min. Samples were spun down at 8000 rpm for 5 min and washed with 1X PBS 3 times. Cells were resuspended in 50 µl of DAPI mounting media (VECTASHIELD® Antifade Mounting Medium with DAPI H-1200-10) and incubated at room temperature for 10 min, away from direct light. The samples were imaged by using a Nikon Ni-E upright microscope equipped with a Yokogawa CSU-W1 spinning-disk head, an Andor iXon 897U EMCCD camera, Nikon Elements AR software, a 60x oil immersion objective, and a 358 nm laser. 15 z-stacks with a thickness of 0.3 µm were collected per image.

### Adaptation and Auxin Plating Assays

We performed adaptation assays as previously described (Eapen et al. 2012; Lee et al. 1998). Cells grown in YEP-Lac overnight were diluted 20-fold in sterile water and plated on a YEP-agar plate containing 2 % galactose. Using micromanipulation, 50 G_1_ cells were isolated and positioned in a grid followed by incubation at 30° C. To quantify the percentage of adapted cells, the number of cells that re-entered cell cycle, grew to a microcolony (3+ cells) after 24 h was divided by the total number of cells. For an auxin plating assays, damage was induced in a YEP-Lac liquid culture by adding galactose at a final concentration of 2 %, as described above. Cells were then transferred onto YEP-agar plates containing 2 % galactose and 1mM IAA or 1 µM 5-Ph-IAA 4 h or 15 h after adding galactose. For each timepoint, >250 cells were scored and categorized as described above for the adaptation assay.

### Quantification and Data Analysis

Graphs were prepared using GraphPad Prism 10 (Dotmatics). Statistical analysis for differences in the percentage of large budded (G_2_/M arrested) cells at different timepoints listed in **S. Table 1** was done using a one-way Anova test in GraphPad Prism 10. Protein quantification of Ddc2-myc blots was done using ImageLab 6.1 (BioRad). To categorize DAPI stained cells based on their morphology and number of DAPI signals, images were captured as described above and viewed using ImageJ with the Fiji addon.

## Acknowledgments

We thank Helle Ulrich for the AID and TIR1 plasmid and Masato Kanemaki for the TIR1(F74G) plasmid. This research was supported by NIH grant R35 GM127029. D.P.W. and S.C.-P. were supported by NIH Genetics Training Grant TM32GM007122. G.M. was supported by NIH Ruth L. Kirschstein Postdoctoral Individual National Research Award F32-GM145156. V.V.E. was supported by a Howard Hughes Medical Institute International Predoctoral Fellowship. We also thank Nikita Alimov, Jessie Ang, and Astré Bouchier for their assistance.

## Author Contributions

Conceptualization and initial data collection was conducted by David P. Waterman Vinay V. Eapen and James E. Haber. Data collection, strain making, imaging, primer design, and data analysis were performed by Felix Zhou, David P. Waterman, Marissa Ashton, Suhaily Caban-Penix, Gonen Memisoglu, and Vinay Eapen. The manuscript was authored and edited by Felix Zhou, David P. Waterman, Gonen Memisoglu, and James E. Haber.

