## Supplemental tables and figures for "Prolonged Cell Cycle Arrest in Response to DNA damage in Yeast Requires the Maintenance of DNA Damage Signaling and the Spindle Assembly Checkpoint"

**S. Table 1 – Comparison of the percentage of large-budded cells back to baseline levels**

| Figure | Strain | Timepoint Comparison | P value | Significance | Post-hoc Test |
|---|---|---|---|---|---|
| 2A | *DDC2-AID* | 0 vs 5 + IAA | <0.0001 | **** | Sidak |
| | | 0 vs 6 + IAA | <0.0001 | **** | Sidak |
| | | 0 vs 7 + IAA | <0.0001 | **** | Sidak |
| | | 0 vs 8 + IAA | 0.0009 | *** | Sidak |
| | | 0 vs 9 + IAA | 0.054 | ns | Sidak |
| 3D | *DDC2-AID CHK1Δ* | 0 vs 5 + IAA | <0.0001 | **** | Sidak |
| | | 0 vs 6 + IAA | <0.0001 | **** | Sidak |
| | | 0 vs 7 + IAA | 0.10 | ns | Sidak |
| | | 0 vs 8 + IAA | 0.25 | ns | Sidak |
| | | 0 vs 9 + IAA | 0.072 | ns | Sidak |
| 2B | *RAD9-AID* | 0 vs 5 + IAA | <0.0001 | **** | Sidak |
| | | 0 vs 6 + IAA | <0.0001 | **** | Sidak |
| | | 0 vs 7 + IAA | <0.0001 | **** | Sidak |
| | | 0 vs 8 + IAA | 0.0055 | ** | Sidak |
| | | 0 vs 9 + IAA | 1.00 | ns | Sidak |
| 3E | *RAD9-AID CHK1Δ* | 0 vs 5 + IAA | 0.00050 | *** | Sidak |
| | | 0 vs 6 + IAA | 0.0052 | ** | Sidak |
| | | 0 vs 7 + IAA | 0.21 | ns | Sidak |
| 2C | *RAD24-AID* | 0 vs 5 + IAA | <0.0001 | **** | Sidak |
| | | 0 vs 6 + IAA | <0.0001 | **** | Sidak |

| | | | | | |
|---|---|---|---|---|---|
| | | 0 vs 7 + IAA | 0.00 | *** | Sidak |
| | | 0 vs 8 + IAA | 0.36 | ns | Sidak |
| | | 0 vs 9 + IAA | 0.90 | ns | Sidak |
| 3F | *RAD24-AID CHK1Δ* | 0 vs 5 + IAA | 0.00010 | *** | Sidak |
| | | 0 vs 6 + IAA | 0.00020 | *** | Sidak |
| | | 0 vs 7 + IAA | 0.10 | ns | Sidak |
| 5D | *RAD53-AID TIR1(F74G)* | 18 vs 18 + IAA | 0.35 | ns | Sidak |
| | | 21 vs 21 + IAA | 0.96 | ns | Sidak |
| | | 24 vs 24 + IAA | 0.42 | ns | Sidak |
| 5E | *RAD9-AID pRAD9-AID* | 18 vs 18 + IAA | 0.80 | ns | Sidak |
| | | 21 vs 21 + IAA | 0.99 | ns | Sidak |
| | | 24 vs 24 + IAA | 0.84 | ns | Sidak |
| S8A | *DDC2-AID MAD2-AID AND MAD2-AID* | 18 + IAA vs 18 + IAA | 0.95 | ns | Sidak |
| | | 21 + IAA vs 21 + IAA | 0.64 | ns | Sidak |
| | | 24 + IAA vs 24 + IAA | 0.97 | ns | Sidak |

**a.** Timepoints are relative to when galactose was added.

**b.** A one-way Anova test was used to test for significant differences.

**S. Table 2 – Strains**

| Strain | Parent | Genotype | Reference |
|---|---|---|---|
| JKM179 | | *MATα ade1 leu2-3 lys5 trp1::hisG ura3-52 hoΔ hmlΔ::ADE1 hmrΔ::ADE1 ade3::GAL::HO* | Lee et al. Cell 1998 |
| DW417 | JKM179 | *HOcse6::HPH TIR1-myc6::URA3* | This study |
| DW418 | DW417 | *Ddc2-AID*-9xMyc::KAN* | This study |
| DW419 | DW417 | *Rad9-AID*-9xMyc::KAN* | This study |
| DW420 | DW417 | *Rad24-AID*-9xMyc::KAN* | This study |
| DW421 | DW417 | *Rad53-AID*-9xMyc::KAN* | This study |
| DW426 | DW417 | *chk1Δ::NAT* | This study |
| DW647 | DW418 | *Ddc2-AID*-9xMyc::KAN chk1Δ::NAT* | This study |
| DW427 | DW419 | *Rad9-AID*-9xMyc::KAN chk1Δ::NAT* | This study |
| DW428 | DW420 | *Rad24-AID*-9xMyc::KAN chk1Δ::NAT* | This study |
| DW429 | DW421 | *Rad53-AID*-9xMyc::KAN chk1Δ::NAT* | This study |
| DW625 | DW417 | *dun1Δ::KAN* | This study |
| DW626 | DW417 | *Dun1-AID*-9xMyc::KAN* | This study |
| DW641 | DW626 | *Dun1-AID*-9xMyc::KAN chk1Δ::NAT* | This study |
| FZ009 | DW417 | *Mad2*-9xMyc-AID::NAT* | This study |
| FZ010 | DW418 | *Ddc2-AID*-9xMyc::KAN Mad2*-9xMyc-AID::NAT* | This study |
| DW455 | DW417 | *mad2Δ::KAN* | This study |
| GM180 | JKM179 | *pGal::Ddc2::LEU2* | (Memisoglu et al. 2019) |
| DW648 | GM180 | *HOcse6::HPH pGal::Ddc2::LEU2* | This study |
| DW649 | DW648 | *HOcse6::HPH pGal::Ddc2::LEU2 mad2Δ::NAT* | This study |

| | | | |
|---|---|---|---|
| DW642 | JKM179 | *HOcse6::HPH Ddc2-AID*-9xMyc::KAN* | This study **a** |
| DW643 | JKM179 | *HOcse6::HPH Rad9-AID*-9xMyc::KAN* | This study **b** |
| DW644 | JKM179 | *HOcse6::HPH Rad24-AID*-9xMyc::KAN* | This study |
| DW645 | JKM179 | *HOcse6::HPH Rad53-AID*-9xMyc::KAN* | This study |
| DW650 | GM180 | *pGal::Ddc2::LEU2 mad2Δ::NAT* | This study |
| GM539 | JKM179 | *Ddc2-9xMyc::KAN* | (Memisoglu et al. 2019) |
| FZ001 | JKM179 | *MATα HOcse6::HPH* | This study |
| JY542 | JKM179 | *HOcse6::HPH tel1Δ::KAN* | This study |
| FZ024 | FZ001 | *HOcse6::HPH Rad9-AID*-9xMyc::KAN* | This study |
| FZ025 | FZ001 | *HOcse6::HPH Rad24-AID*-9xMyc::KAN* | This study |
| FZ026 | FZ001 | *HOcse6::HPH Rad53-AID*-9xMyc::KAN* | This study |
| FZ173 | FZ025 | *HOcse6::HPH Rad24-AID*-9xMyc::KAN TIR1(F74G)::URA3* | This study |
| FZ174 | FZ024 | *HOcse6::HPH Rad9-AID*-9xMyc::KAN TIR1(F74G)::URA3* | This study |
| FZ175 | FZ026 | *HOcse6::HPH Rad53-AID*-9xMyc::KAN TIR1(F74G)::URA3* | This study |
| YSL53 | JKM179 | *HOcse5::URA3* | (Lee et al. 1998) **c** |
| GEM188 | JKM179 | *HOcse2::LYS2* | This study |
| FZ201 | DW419 | *Rad9-AID*-9xMyc::KAN pRAD9-AID*-9xMyc* | This study |
| FZ155 | DW417 | *bfa1Δ::KAN* | This study |
| yMA11 | JKM179 | H2A-S129A H2B-T129A | This study |
| yMA12 | JKM179 | H2AS129E H2B-T129E | This study |
| yMA13 | JKM179 | H2B-T129A | This study |
| yMA14 | JKM179 | H2B-T129E | This study |

| | | | |
|---|---|---|---|
| yBL257 | JKM179 | H2A-S129E | This study |
| yBL259 | JKM179 | H2A-S129A | This study |

**a.** *HOcse6::HPH* is the an HO-cut site 42 kb away from the centromere on chromosome VI.
**b.** *HOcse5::URA3* is the an HO-cut site 36 kb away from the centromere on chromosome V.
**c.** *HOcse2::LYS2* is the an HO-cut site 230 kb away from the centromere on chromosome II.

**S. Table 3 – Primers**

| Oligo | Sequence | Use |
|---|---|---|
| GAT1p1B | GCTCAGTGTGCGTTATGCTT | Primer to add the second HO-cut site on chromosome VI |
| GAT1p2B | TTCAGGTCTCGGTTGCTCTT | Primer to add the second HO-cut site on chromosome VI |
| VE162 Ddc2-AID For | ATCTAACCACACTAGAGGAGGCCGATTCATTATATATCTCAATGGGACTGCCTAAAGATCCAGCCAAACCTCC | C-terminal AID tag for Ddc2 (forward) |
| VE163 Ddc2-AID Rev | ATTACAAGGTTTCTATAAAGCGTTGACATTTTCCCCTTTTGATTGTTGCCCAGTATAGCGACCAGCATTCACATAC | C-terminal AID tag for Ddc2 (reverse) |
| DW217 Rad9-AID 1F | GGTTTTCACGATGATATTACGGACAATGATATATACAACACTATTTCTGAGGTTAGACCTAAAGATCCAGCCAAACCTCC | C-terminal AID tag for Rad9 (forward) |
| DW218 Rad9-AID 1R | CTAAATTTTTTTTATTTAATCGTCCCTTTCTATCAATTATGAGTTTATATATTTTTATAATTCAGTATAGCGACCAGCATTCACATAC | C-terminal AID tag for Rad9 (reverse) |
| DW208 Rad24-AID 1F | CAGATTCAGATCTGGAAATACTCCCTAAAGATCCAGCCAAACCTCC | C-terminal AID tag for Rad24 (forward) |
| DW209 Rad24-AID 1R | GTGGAATATTTCCTGGGGTTTCTCGTCAAATTTAAAGAGTAAAAAGCCTAAAGATCCAGCCAAACCTCC | C-terminal AID tag for Rad24 (reverse) |
| DW199 Rad53AID 1F | GGTTAAAAGGGCAAAATTGGACCAAACCTCAAAAGGCCCCGAGAATTTGCAATTTTCGCCTAAAGATCCAGCCAAACCTCC | C-terminal AID tag for Rad53 (forward) |
| DW200 Rad53AID 1R | CCATCTTCTCTCTTAAAAAGGGGCAGCATTTTCTATGGGTATTTGTCCTTGGCAGTATAGCGACCAGCATTCACATAC | C-terminal AID tag for Rad53 (reverse) |
| dw418 Dun1 1F | CGAGAGTAACAAGTAAAGGGGCTTAACATACAGTAAAAAAGGCAATTATAGTGAAGATGCCTTGACAGTCTTGACGTGC | Genomic deletion of Dun1 (forward) |
| dw419 dun1 1R | GATACTTGGAAAAATCCAGATTCAAACAATGTTTTTGAAATAATGCTTCTCATGTTTACGCACTTAACTTCGCATCTG | Genomic deletion of Dun1 (reverse) |
| Sp01Dun1 aidF | CAATAAAATACCCAAAACATACTCAGAATTATCTTGCCTCCCTAAAGATCCAGCCAAACCTCC | C-terminal AID tag for Dun1 (forward) |
| SP02Dun1 aidR | CCAGATTCAAACAATGTTTTTGAAATAATGCTTCTCATGTCAGTATAGCGACCAGCATTCACATAC | C-terminal AID tag for Dun1 (reverse) |
| BL189-Chk1 1F | TCAGCCACTGGTCATCCCGT | Genomic deletion Chk1 (forward) |

| | | |
|---|---|---|
| BL190-Chk1 1R | GTTGGGGGGAGATGGTAACG | Genomic deletion Chk1 (reverse) |
| FZ013 Mad2-AID 1F | CATTCTCTACCAACGATCATAAAGTTGGT GCGCAGGTCAGCTATAAATATCCTAAAGA TCCAGCCAAACCTCC | C-terminal AID tag for Mad2 (forward) |
| FZ014 Mad2-AID 1R | CGAGATTTTTTGGACTTCCGTCTTTTTTT TTTTTTTTGACTTGAATTCTAGATATCATC GATGAATTCGAGCTCG | C-terminal AID tag for Mad2 (reverse) |
| FZ096 Mad1-AID 1F | GGCAACAATAACATTGCGTCTGTGGGAAC AGCGACAAGCCAAA CCTAAAGATCCAGCCAAACCTCC | C-terminal AID tag for Mad1 (forward) |
| FZ097 Mad1-AID 1R | GGAGTTTATCATATTATAAAACCGATTAC TATTATCTATTAGAAATGTATATACAC GATATCATCGATGAATTCGAGCTCG | C-terminal AID tag for Mad1 (reverse) |
| FZ147 Bub2-AID 1F | GACCACTTGACCGACCCAGACATATATAT ACCG CGTACGCTGCAGGTCGAC | C-terminal AID tag for Bub2 (forward) |
| FZ148 Bub2-AID 1R | CGTTGTAGAATTAAACGATAAAATATAAT ATTTCTTCACATAGT ATCGATGAATTCGAGCTCG | C-terminal AID tag for Bub2 (reverse) |
| FZ149 Bfa1-AID 1F | CCTATATGTATGAAATCAGGAACATGGTA ATCAATTCGACAAAAGAT CGTACGCTGCAGGTCGAC | C-terminal AID tag for Bfa1 (forward) |
| FZ150 Bfa1-AID 1R | CTCAAGATAACGGTAAAGAAACAGTTATA AGAAGGCTAAAGGG ATCGATGAATTCGAGCTCG | C-terminal AID tag for Bfa1 (reverse) |
| FZ152 Bfa1 2F | GATGTTTTCGGAGACAATTTGGTTACTG | Genomic deletion Bfa1 (forward) |
| FZ153 Bfa1 2R | GACGCGAAATGTCGGCG | Genomic deletion Bfa1 (reverse) |
| GM474 | CGGGCCGTTTAGCGGAATGGATGAGTAAG TATGGTTGCACGATCTAAATAAATTCGTT TTCAATGATTAAAATAGCATAGTCGGGTT TTTCTTTTAGTTTCAGCTTTCCGCAACAGT ATAATTTTATAAACCCTGGTTTTGGTTTTG TAGAGTGGTTTTGTGAACAACTGGTTTGT TGAAAAAGATCACTGGAATTA | Donor sequence template to insert an HO-cut site at *LYS2*. Use with plasmids bG059 or bG060. Red is the insertion for the HO-cut site. |
| FZ178 Lys2 gRNA 1 | ACCCATTTAACACCTGCCAT | gRNA for *LYS2*. Inserted into plasmid bRA90. |
| FZ179 Lys2 gRNA 2 | GGTTTGGCCGAAGGTTATAG | gRNA for *LYS2*. Inserted into plasmid bRA90. |
| FZ180 Rad9 1F | CGTACGCTGCAGGTCGAC | Forward primer to add AID tag to Rad9 in plasmid pFL36.1. Use with primer |

| | | |
|---|---|---|
| | | FZ181 and plasmid pJH2892. |
| FZ181 Rad91R | GGTGGTggcgcgccTTTTTAGCTAGT | Forward primer to add AID tag to Rad9 in plasmid pFL36.1. Use with primer FZ180 and plasmid pJH2892. Lower case is the location of a restriction enzyme site AscI. Red is a random sequence to assist with cutting by AscI. |
| FZ152 Bfa1 2F | GATGTTTTCGGAGACAATTGGTTACTG | FW primer to check for Bfa1 deletion. 326bp upstream of start codon. Use with yeast haploid deletion collection. |
| FZ153 Bfa1 2R | GACGCGAAATGTCGGCG | Rev primer to check for Bfa1 deletion. 101bp downstream of stop codon. Use with yeast haploid deletion collection. |
| HTA1 gRNA1F | AACGTTACCATTGCCCAAGGgtttt | Guide RNA to cut HTA1 using CRISPR/Cas9. |
| HTA2 gRNA 1F | AATGTTACCATCGCCCAAGGgtttt | Guide RNA to cut HTA2 using CRISPR/Cas9. |
| HTB1 T129 gRNA | AGTACTCTTCCTCTACTCAAgatca | Guide RNA to cut HTB1 using CRISPR/Cas9. |
| HTB2 T129 gRNA | AATACTCCTCCTCTACTCAAgatca | Guide RNA to cut HTB2 using CRISPR/Cas9. |
| HTB1 T129A 80 mer | CTCTGAAGGTACTAGAGCTGTTACgAAGT ACTCgTCCTCTGCTCAAGCATAATGAAATC ACTTCCTTTGGTTATAATTAATAT | Repair template to replace threonine at 129 on HTB1 with alanine. |
| HTB1 T129E 80 mer | CTCTGAAGGTACTAGAGCTGTTACgAAGT ACTCgTCCTCTGAACAAGCATAATGAAAT CACTTCCTTTGGTTATAATTAATAT | Repair template to replace threonine at 129 on HTB1 with glutamic acid. |
| HTB2 T129E 80 mer | CTCCGAAGGTACTAGGGCTGTTACgAAAT ACTCgTCCTCTGAACAAGCCTAAGTCACTC ACTAGGTATTGTGATTTAGTCATG | Repair template to replace threonine at 129 on HTB2 with glutamic acid. |
| HTB2 T129A 80 mer | CTCCGAAGGTACTAGGGCTGTTACgAAAT ACTCgTCCTCTGCTCAAGCCTAAGTCACTC ACTAGGTATTGTGATTTAGTCATG | Repair template to replace threonine at 129 on HTB2 with alanine. |
| HTA1 S129A 80mer | AAGGTGGTGTTTTGCCAAACATCCATCAA AACTTGTTGCCAAAGAAGTCTGCCAAGGC TACCAAGGCTgctCAAGAATTA | Repair template to replace serine at 129 on HTA1 with alanine. |
| HTA1 S129E 80mer | AAGGTGGTGTTTTGCCAAACATCCATCAA AACTTGTTGCCAAAGAAGTCTGCCAAGGC TACCAAGGCTgaaCAAGAATTA | Repair template to replace serine at 129 on HTA1 with glutamic acid |

| | | |
|---|---|---|
| HTA2 S129A 80mer | CCAAGGTGGTGTTTTGCCAAACATTCACC AAAACTTGTTGCCAAAGAAGTCTGCCAAG ACTGCCAAAGCTgctCAAGAAC | Repair template to replace serine at 129 on HTA2 with alanine. |
| HTA2 S129E 80mer | CCAAGGTGGTGTTTTGCCAAACATTCACC AAAACTTGTTGCCAAAGAAGTCTGCCAAG ACTGCCAAAGCTgaaCAAGAAC | Repair template to replace serine at 129 on HTA2 with glutamic acid. |

**S. Table 4 – Plasmids**

| Name | Plasmids | Backbone | Reference |
|---|---|---|---|
| pKan-9xMyc-AID | pJH2892 | pSM409 | (Morawska and Ulrich 2013) |
| pNAT-9xMyc-AID | pJH2899 | pSM409 | (Morawska and Ulrich 2013) |
| osTIR1::*URA3* | pNHK53 | | (Nishimura et al. 2009) |
| GAL-DDC2 | pML100 | pML95 | (Paciotti et al. 2000) |
| ADH1-OsTIR1(F74G) | pMK420 | | (Yesbolatova et al. 2020) |
| bRA90 | bRA90 | | (Anand et al. 2017) |
| bG059 | bG059 | bRA90 | This study |
| bG060 | bG060 | bRA90 | This study |
| pRad9-3HA | pFL36.1 | pRS306 | (Lazzaro et al. 2008) |
| pRad9-9xMyc-AID | pFZ052 | pRS306 | This study |
| pBL15 – HTA1 gRNA1 | pBL15 | BRA89 | This study |
| pBL16 – HTA2 gRNA2 | pBL16 | BRA89 | This study |
| pKL004 – HTB1 gRNA1 | pKL004 | BRA89 | This study |
| pKL005 – HTB2 gRNA1 | pKL005 | BRA89 | This study |

### S. Table 5 – Key Resources

| Reagent or Resource | Source | Identifier |
|---|---|---|
| **Antibodies** | | |
| Mouse monoclonal anti-Rad53 | AbCam | Cat # ab166859; RRID: AB_2801547 |
| Rabbit polyclonal anti-Rad53 | AbCam | ab104232 |
| Mouse monoclonal anti-Myc | AbCam | Cat# ab16918; RRID: AB_30256 |
| Mouse monoclonal anti-Pgk1 | AbCam | Cat# ab32, RRID: AB_30359 |
| Rabbit polyclonal anti-Rad9 | (Usui, Foster, and Petrini 2009) | N/A |
| ECL TH Anti-mouuse IgG horseradish peroxidase from sheep | GE Healthcare | NXA931V Lot 16937010 |
| ECL TH Anti-rabbit IgG horseradish peroxidase from donkey | GE Healthcare | NA934V lot 6969611 |
| **Chemicals, Peptides, and Recombinant Proteins** | | |
| Indole-3-acetic acid | Sigma-Aldrich | I3750-25G-A |
| 5-Ph-IAA | Sigma-Aldrich | SML3574-25MG |
| Formaldehyde | Sigma-Aldrich | 47608 |
| VECTASHIELD® Antifade Mounting Medium with DAPI | Vector-Laboratories | Cat. No. H-1200 |
| Prometheus Protein Biology Products 20-313 OneBlock™ Western-CL Blocking Buffer, For Chemiluminescent Blots | Genesee Scientific | Cat #: 20-313 |
| ECL™ Prime Western Blotting System | Millipore Sigma | GERPN2232 |
| **Experimental Models: Organisms/Strains** | | |
| S. cerevisiae: Strain background S228c. Strains are listed in S. Table 2 | This paper | N/A |
| **Oligonucleotides** | | |
| Oligonucleotides tagging proteins with an AID tag and deleting genes are listed in S. Table 3 | This paper | N/A |
| **Recombinant DNA** | | |
| Plasmids used in this study are listed in S. Table 4. | This paper | N/A |
| **Software and Algorithms** | | |
| Prism 7.00 | GraphPad Software, Inc. | N/A |
| Image Lab | Bio-Rad | N/A |
| FiJi | | N/A |

### Supplemental Figures

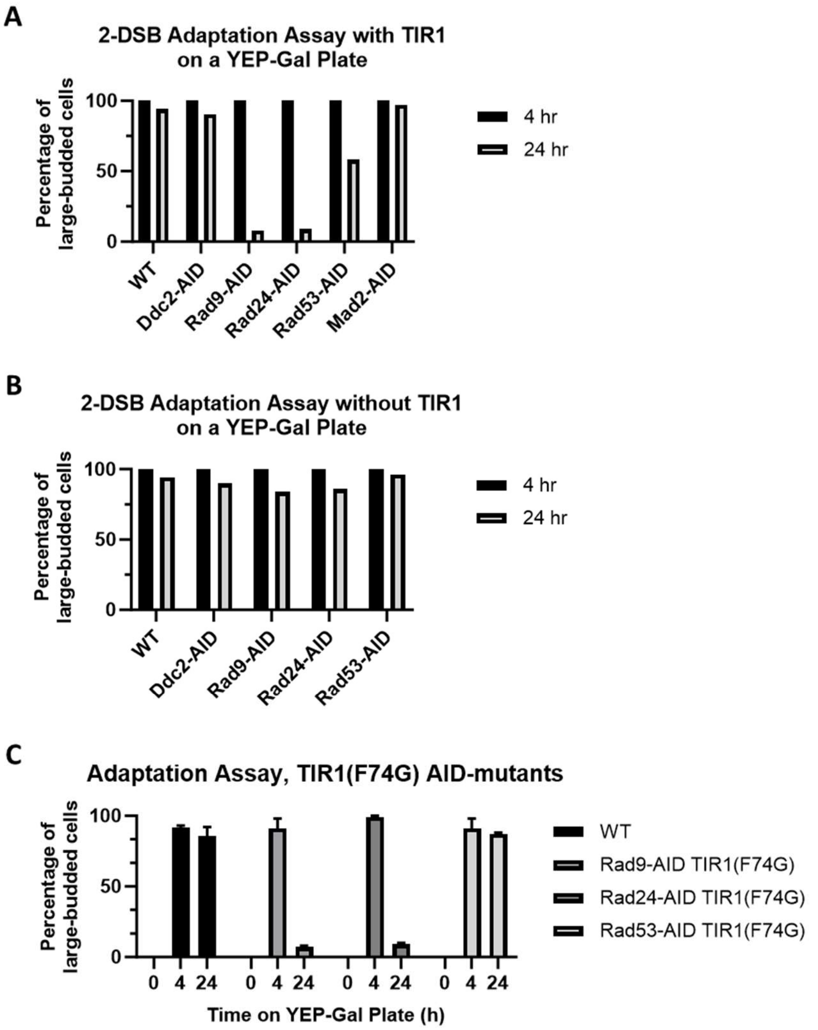

**Figure S1. Adaptation assay of AID-tagged checkpoint activation proteins**
(A) Adaptation assay of 50 $G_1$ cells on a YEP-Gal plate after 24 h for 2-DSB (WT), 2-DSB *DDC2-AID*, 2-DSB *RAD9-AID*, 2-DSB *RAD24-AID*, and 2-DSB *RAD53-AID* with TIR1. (B) Adaptation assay of 50 $G_1$ cells on a YEP-Gal plate after 24 h for 2-DSB (WT), 2-DSB *DDC2-AID*, 2-DSB *RAD9-AID*, 2-DSB *RAD24-AID*, and 2-DSB *RAD53-AID* without TIR1. (C) Adaptation assay of 50 $G_1$ cells on YEP-Gal plate after 24 h for 2-DSB (WT), 2-DSB *RAD9-AID* TIR1 (F74G), 2-DSB *RAD24-AID* (F74G), and 2-DSB *RAD53-AID* TIR1(F74G).

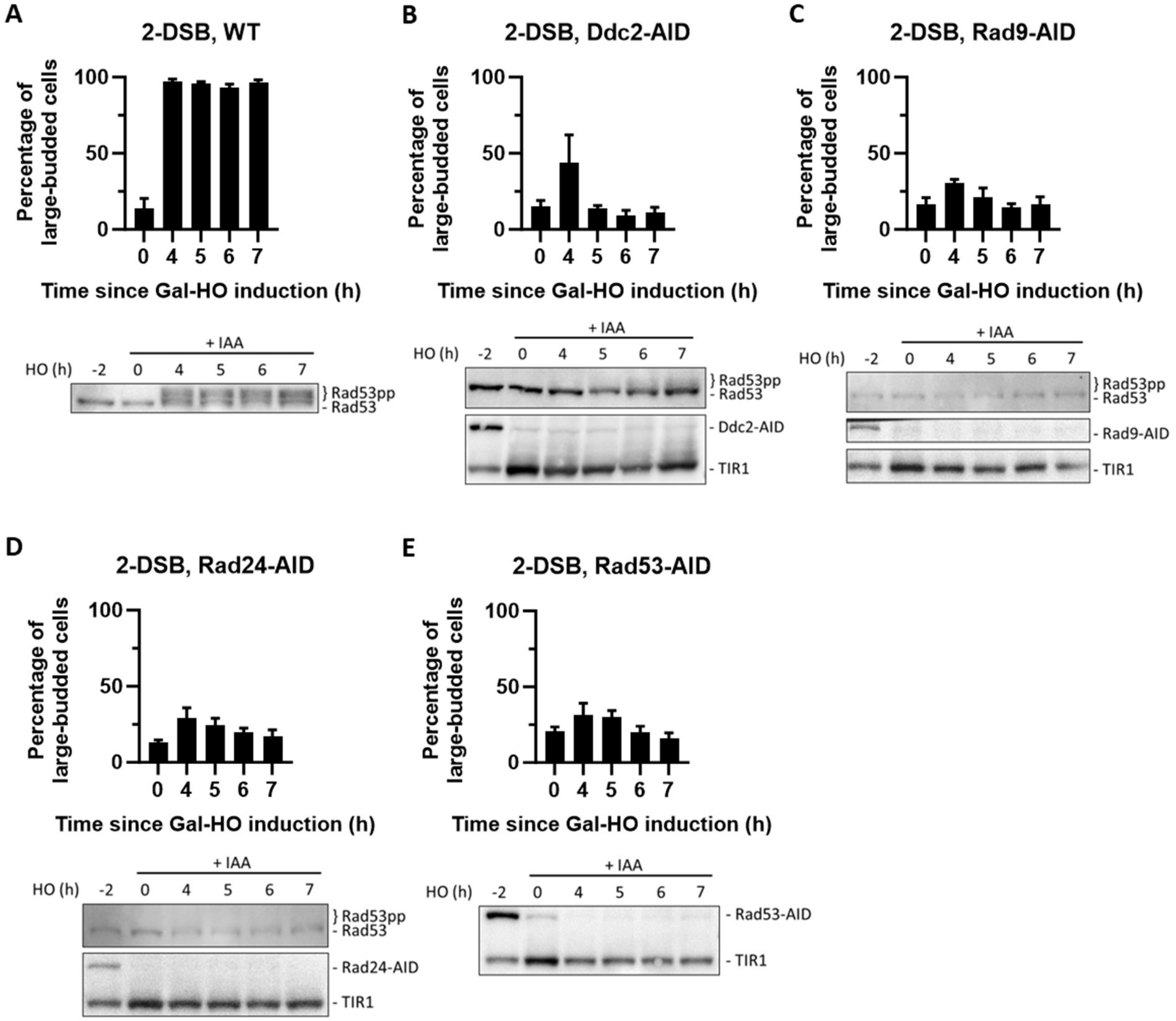

**Figure S2. AID-tagged checkpoint proteins readily degrade with auxin**

(A) Morphological profile of a 2-DSB background where 1 mM Auxin (IAA) was added 2 h before DSB-induction with galactose. Western blot of a 2-DSB strain probed with α-Rad53. α-Rad53 shows both an unphosphorylated protein and multiple phosphorylated species. (B) Morphological profile of *DDC2-AID* in a 2-DSB background where IAA was added 2 h before galactose. Western blot probed with α-Rad53 and α-Myc. α-Rad53 shows both an unphosphorylated protein and multiple phosphorylated species. α-Myc shows Ddc2-AID degradation and TIR1-Myc as a loading control. (C) Same as (B) for 2-DSB *RAD9-AID*. α-Myc probe shows Rad9-AID degradation with IAA. (D) Same as (B) for 2-DSB *RAD24-AID*. α-Myc probe shows Rad24-AID degradation with IAA. (E) Same as (B) for 2-DSB *RAD53-AID*. α-Myc probe shows Rad53-AID degradation with IAA.

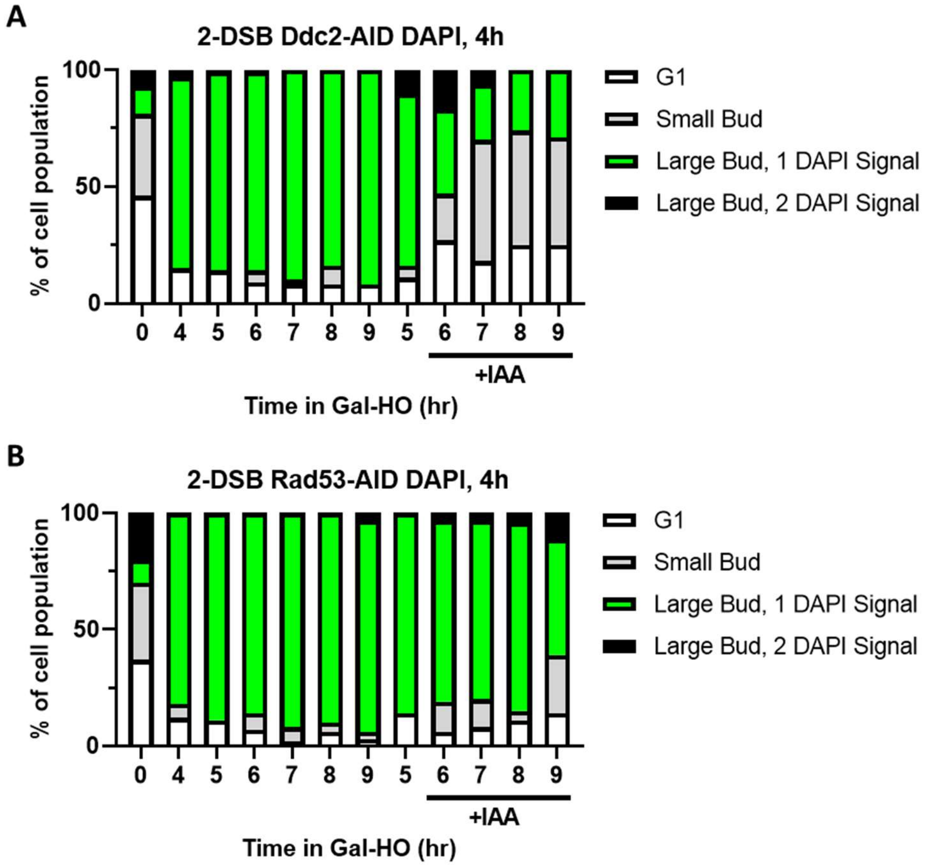

**Figure S3. Cell cycle profile as determined by budding and DAPI staining in Ddc2-AID and Rad53-AID mutants ± IAA 4 h after galactose**

(A) Profile of DAPI stained cells in a 2-DSB *DDC2-AID* strain after HO induction. Cultures were split 4 h after Gal-HO induction. Cells were divided based on cell morphology and number of DAPI signals. (B) Same as (A) for 2-DSB *RAD53-AID*.

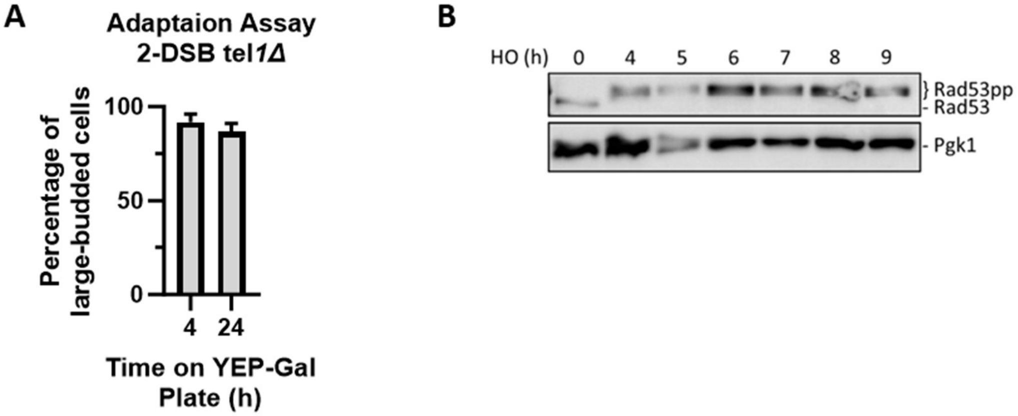

**Figure S4. Tel1 is not required for DDC activation or Rad53 phosphorylation**
(A) Adaptation assay for a *tel1*Δ strain. Cultures were grown in YEP-Lac and put on a YEP-Gal plate. 50 $G_1$ cells were selected to monitor their morphology after 4 and 24 h on the YEP-Gal plate. (B) Western blot probing α-Rad53 in a 2-DSB *tel1*Δ. α-Pgk1 probed as a loading control.

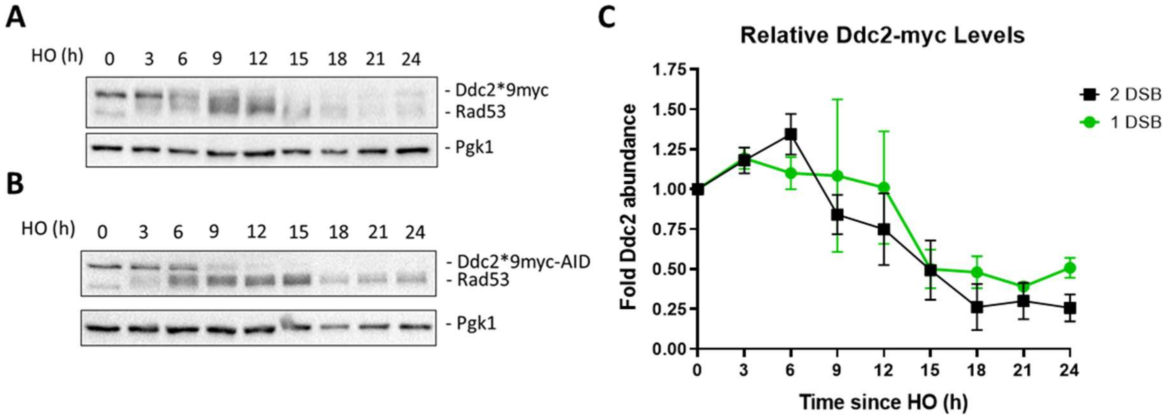

**Figure S5. Relative levels of Ddc2 levels decrease after DSB induction**
(A-B) Western blots probed with α-Myc for Ddc2-9xMyc and Ddc2-9xMyc-AID in a 1-DSB and 2-DSB, respectively. α-Pgk1 is used as a loading control. (C) Relative levels of Ddc2 in a 1-DSB and 2-DSB strain up to 24 h after DSB induction.

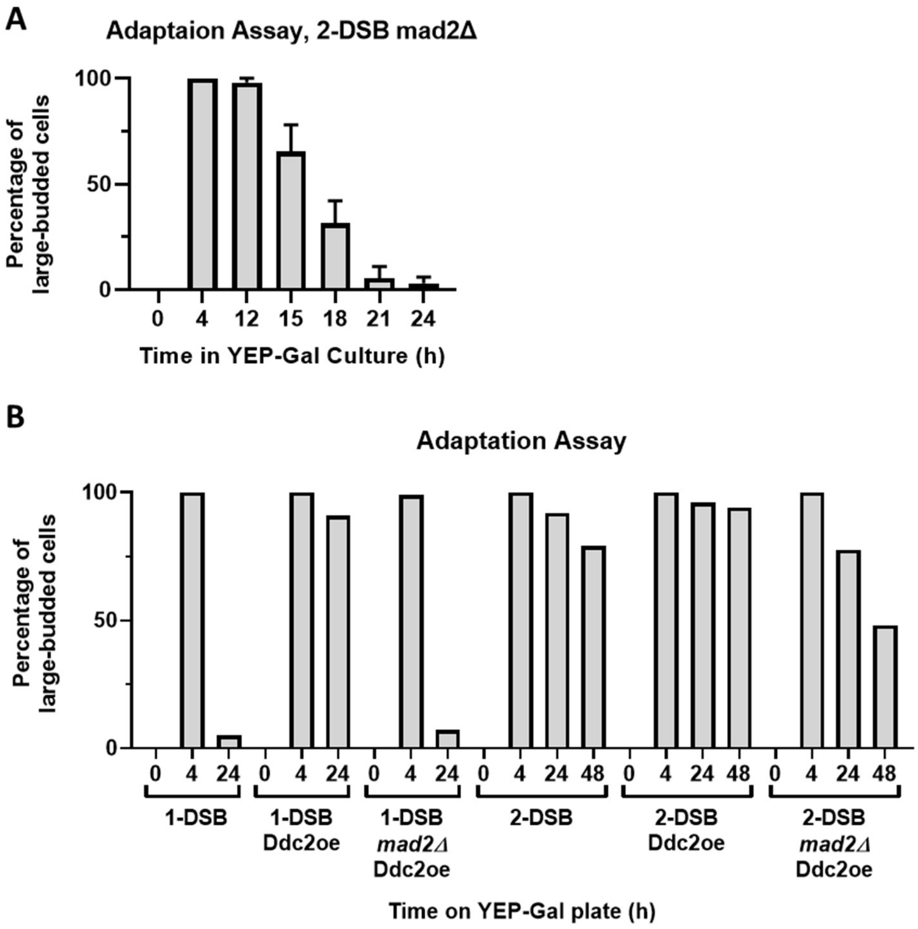

**Figure S6. Mad2 is required for permanent arrest in a 2-DSB strain**
(A) Adaptation assay of 1-DSB and 2-DSB where morphology is measured for up to 24 h and 48 h, respectively. A second copy of Ddc2 with a GAL1,10 promotor, Ddc2 overexpression (Ddc2oe), was integrated into the 1-DSB and 2-DSB strains. *MAD2* was deleted in both backgrounds with the second copy of Ddc2. (B) Adaptation assay for a *mad2*Δ in a 2-DSB strain.

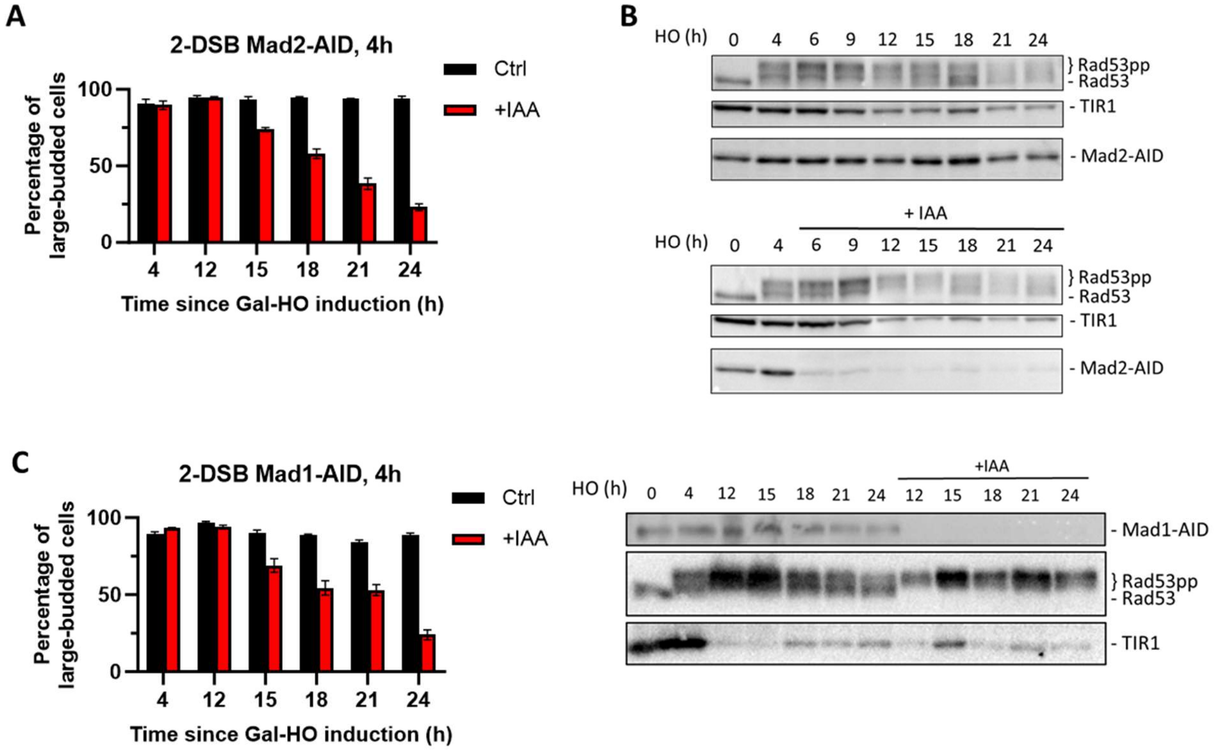

**Figure S7. Mad1 and Mad2 are required for permanent arrest in a 2-DSB strain**
(A) Morphological profile of *MAD2-AID* after HO induction on a YEP-Gal or YEP-Gal-IAA plate. Galactose was added to an overnight culture of Mad2-AID in YEP-Lac. 4 h after adding galactose, cells were added to a YEP-Gal plate or a YEP-Gal-IAA (1mM IAA) plate. The morphology of cells was measured for up to 24 h on each plate. (B) Western blot probed with α-Rad53 and α-Myc. α-Rad53 shows both an unphosphorylated protein and multiple phosphorylated species. α-Myc shows Mad2-AID degradation and TIR1-Myc as a loading control. The top western blot samples were treated with ethanol and bottom western blot samples were treated with IAA. (C) Same as (A) for *MAD1-AID*. Western blot probed with α-Rad53 and α-Myc. α-Rad53 shows both an unphosphorylated protein and multiple phosphorylated species. α-Myc shows Mad1-AID degradation and TIR1-Myc as a loading control.

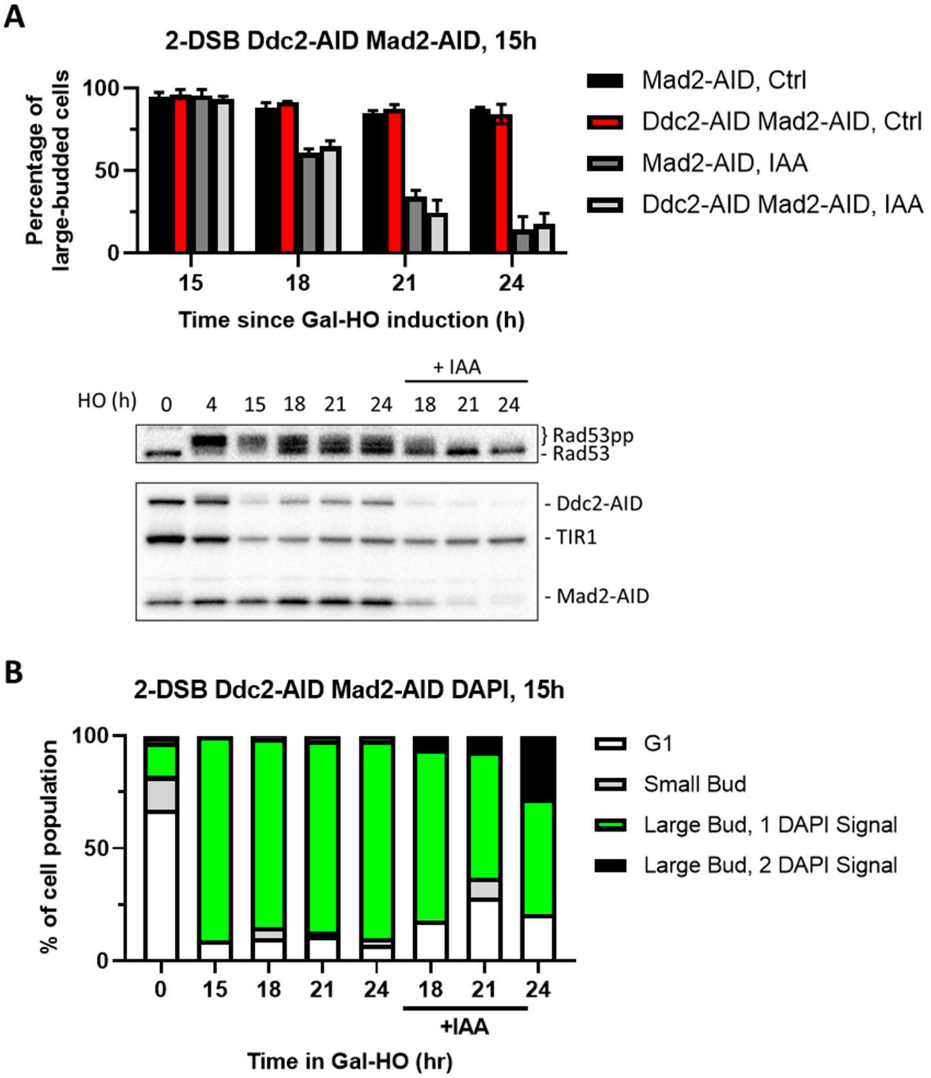

**Figure S8. Degradation of Ddc2 and Mad2 at 15 h releases cells from checkpoint arrest**
(A) Morphological profile of a 2-DSB *MAD2-AID* and *DDC2-AID MAD2-AID* strains after HO induction. Cultures were added onto YEP-Gal ± IAA plates 15 h after adding HO induction. Ctrl samples were plated on a YEP-Gal plate and IAA samples were plated on a YEP-GAL-IAA plate. Western blot probed with α-Rad53 and α-Myc. α-Rad53 shows both an unphosphorylated protein and multiple phosphorylated species. α-Myc shows Ddc2-AID Mad2-AID degradation and TIR1-Myc as a loading control. (B) Profile of DAPI stained cells in a 2-DSB *DDC2-AID MAD2-AID* strain after HO induction. Cultures were split 15 h after HO induction and treated with either IAA or ethanol. Cells were divided based on cell morphology and number of DAPI signals.

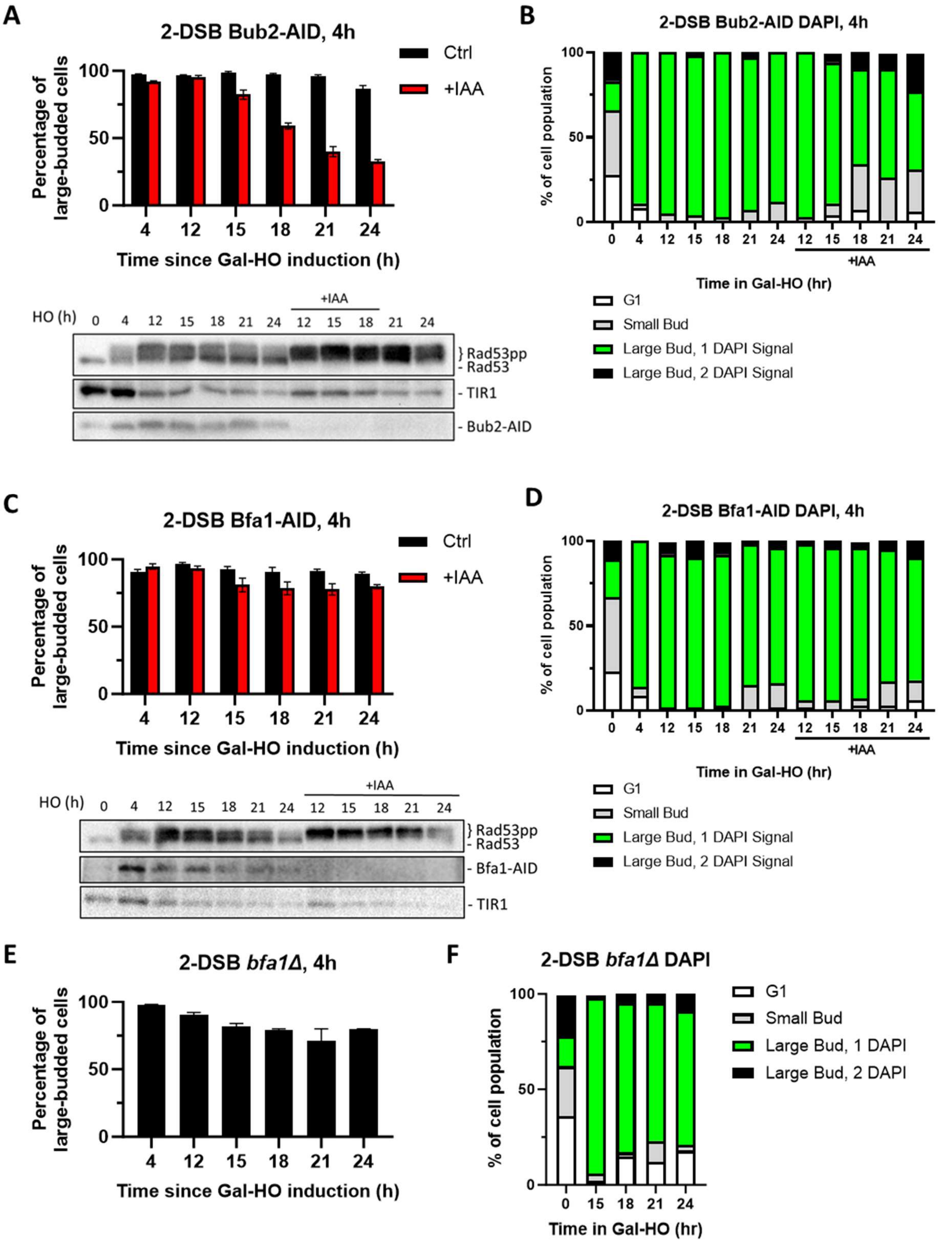

**Figure S9. Bub2 but not Bfa1 is required for prolonged arrest**

(A) Morphological profile of a 2-DSB *BUB2-AID* strain with the auxin-Gal plating assay. Cultures were added onto YEP-Gal ±IAA plates 4 h after HO induction. Western blot probed with α-Rad53 and α-Myc. α-Rad53 shows both an unphosphorylated protein and multiple phosphorylated species. α-Myc shows Bub2-AID degradation and TIR1-Myc as a loading control. (B) Profile of DAPI stained cells in a 2-DSB *BUB2-AID* strain after HO induction. Cultures were split 15 h after HO induction and treated with either IAA or ethanol. Cells were divided based on cell morphology and number of DAPI signals. (C) Same as (A) for a 2-DSB *BFA1-AID* strain. (D) Same as (B) for a 2-DSB *BFA1-AID* strain. (E) Same as (A) for a 2-DSB *bfa1*Δ strain. (F) Same as (B) for 2-DSB *bfa1*Δ strain

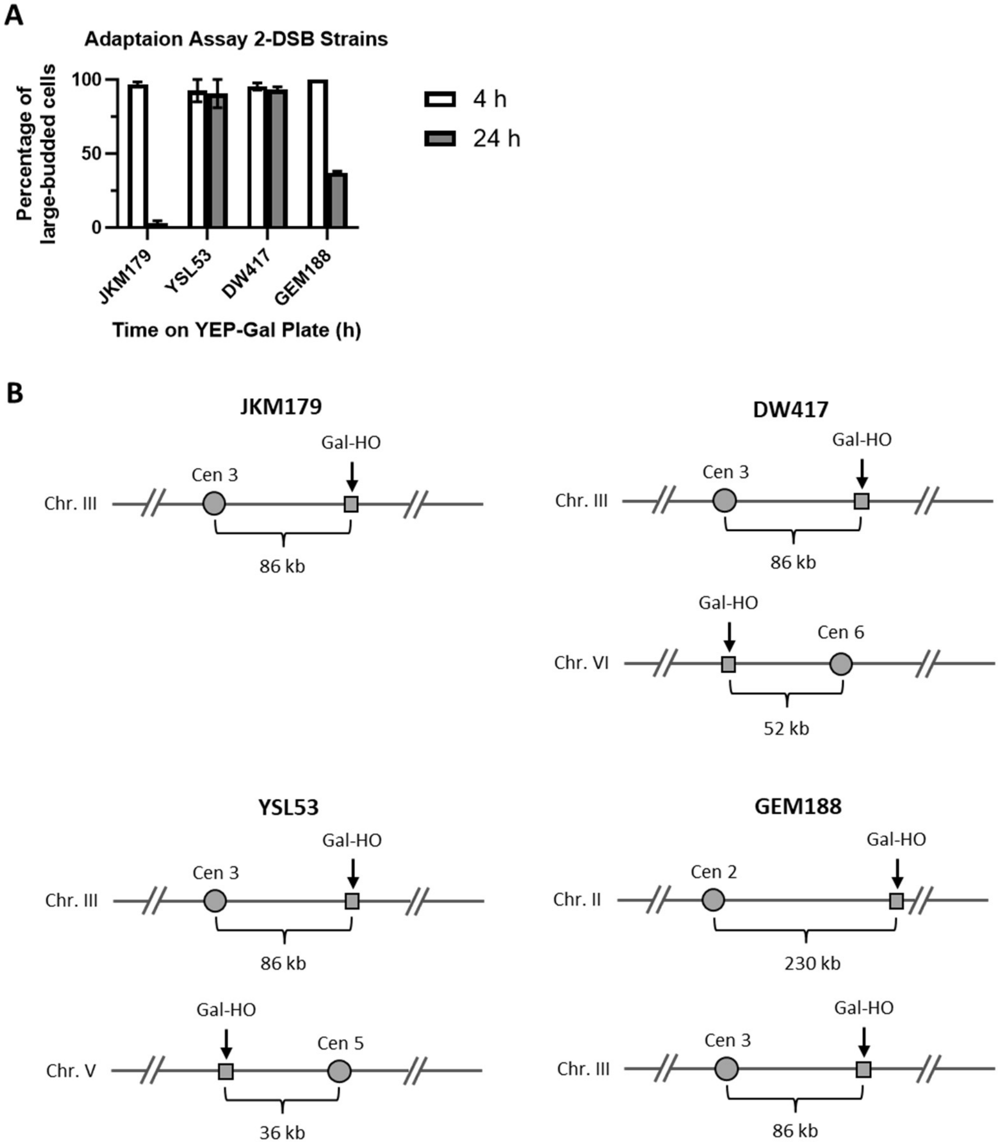

**Figure S10. Adaptation assay of different 2-DSB strains**

(A) Adaptation assay of 1-DSB and 2-DSB strains tracking the morphology of 50 $G_1$ cells on a YEP-Gal plate. The percentage of $G_2$/M arrested cells was shown 4 h and 24 h after placement of YEP-Gal plates. JKM179 is a 1-DSB strain with an HO-cut site in the *MAT* locus on chromosome III 86 kb away from the centromere. DW417 is a 2-DSB strain derived from JKM179 with an additional HO-cut site on chromosome VI 52 kb away from the centromere. YSL53 is a 2-DSB strain derived from JKM179 with an additional HO-cut site at the *URA3* locus on chromosome V 36 kb away from the centromere (Lee et al. 1998). GEM188 is a 2-DSB strain derived from JKM179 with an additional HO-cut site at *LYS2* on chromosome II 230 kb away from the centromere. (B) Cartoon representations of strains showing the location of the HO-cut sites relative to their respective centromeres.

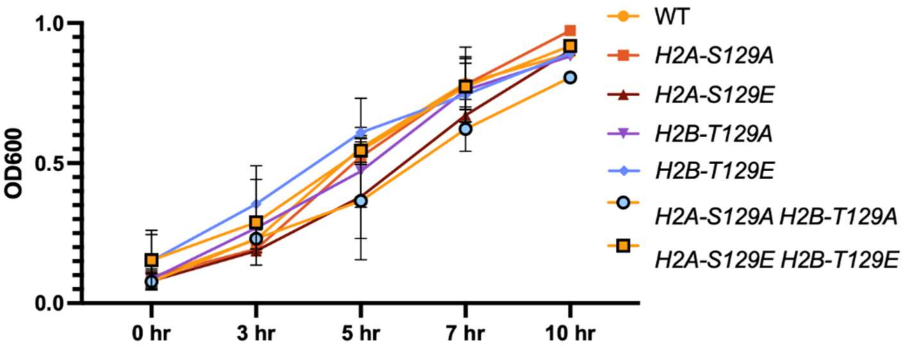

**Figure S11. Phosphomimetic and non-phosphorylatable mutants of histone H2A and H2B do not affect the growth rate of cells.**

Growth rate of strains were measured in YPD (2% dextrose) in H2A and H2B mutants for up to 10 h. Cultures were grown in YPD until they reached an OD600 of 0.1. The OD of each strain was then measured at 3, 5, 7, and 10 h.
